# Mesothelioma Interactome with 367 Novel Protein-Protein Interactions

**DOI:** 10.1101/459065

**Authors:** Kalyani B. Karunakaran, Naveena Yanamala, Gregory Boyce, Madhavi K. Ganapathiraju

## Abstract

Malignant pleural mesothelioma (MPM) is an aggressive cancer of the thorax with a median survival of one year. We constructed an ‘MPM interactome’ with over 300 computationally predicted PPIs and over 1300 known PPIs of 62 literature-curated genes whose activity affects MPM. Known PPIs of the 62 MPM associated genes were derived from BioGRID and HPRD databases. Novel PPIs were predicted by applying the HiPPIP algorithm, which computes features of protein pairs such as cellular localization, molecular function, biological process membership, genomic location of the gene, gene expression in microarray experiments, protein domains and tissue membership, and classifies the pairwise features as *interacting* or *non-interacting* based on a random forest model. To our satisfaction, the interactome is significantly enriched with genes differentially expressed in MPM tumors compared with normal pleura, and with other thoracic tumors. The interactome is also significantly enriched with genes whose high expression has been correlated with unfavorable prognosis in lung cancer, and with genes differentially expressed on crocidolite exposure. 28 of the interactors of MPM proteins are targets of 147 FDA-approved drugs. By comparing differential expression profiles induced by drug to profiles induced by MPM, potentially repurposable drugs are identified from this drug list. Development of PPIs of disease-specific set of genes is a powerful approach with high translational impact – the interactome is a vehicle to piece together an integrated view on how genes associated with MPM through various high throughput studies are functionally linked, leading to clinically translatable results such as clinical trials with repurposed drugs. The PPIs are made available on a webserver, called *Wiki-Pi MPM* at http://severus.dbmi.pitt.edu/wiki-MPM with advanced search capabilities.

**One Sentence Summary:** Mesothelioma Interactome with 367 novel protein-protein interactions may shed light on the mechanisms of cancer genesis and progression

## Introduction

Internal organs such as heart and lung, and body cavities such as thoracic and abdominal cavities, are covered by a thin slippery layer of cells called the “mesothelium”. This protective layer prevents organ adhesion and plays a number of important roles in inflammation and tissue repair (1). The mesothelia that cover the heart, lung and abdominal cavity are specifically called *pericardium, pleura* and *peritoneum*, respectively. Mesothelioma is the cancer that affects these layers. Most types of mesothelioma invade surrounding tissues and blood vessels to form secondary tumors and metastasize to different locations in the body (2). Mesotheliomas of pleura account for ~90% of malignant mesotheliomas and have a short median survival, of only 1 year (3, 4).

Malignant pleural mesothelioma (MPM) is associated with exposure to asbestos over a long period of time (5). The disease onset occurs after a long period after exposure, and this latency makes its detection difficult (2). Only a small fraction of the population exposed to asbestos develops malignant mesothelioma, and the disease tends to cluster in families, which suggests the involvement of a genetic component in the disease (6). Germline mutations were observed in BAP1 gene (7, 8), and inactivating mutations in NF2 gene in pleural mesothelioma (9, 10). Recently, twenty four germline mutations were identified in 13 genes from 198 patients with malignant mesothelioma of pleural or peritoneal origin (11). Non-invasive pre-malignant phase is not seen in mesothelioma unlike other tumors, which necessitates expeditious discovery of genetic predispositions, molecular mechanisms and therapeutics for the disease (12).

60% of the disease-associated missense mutations perturb protein-protein interactions (PPIs) in human genetic disorders (13). PPIs are intricately involved in biological functions and disease mechanisms, and may be exploited for drug-discovery (13). The molecular mechanisms of disease are often revealed by the PPIs of disease-associated genes. For example, the involvement of transcriptional deregulation in the pathogenesis of MPM was identified through mutations detected in BAP1 and its interactions with several proteins including BRCA1 revealed by co-immunoprecipitation (14). The PPI of BRCA1 with BAP1 was also central in understanding its role in growth-control pathways and cancer (15). Studies on BAP1 and BRCA1 later provided the basis for several clinical trials including testing of the drug Vinorelbine as a second line therapy for patients with MPM (16).

Thus, the knowledge of PPIs, if available to biologists specializing in the study of a certain disease, gene or a pathway, would lead to insightful results that advance understanding of disease biology. Despite their importance, only about 10-15% of expected PPIs in the human protein interactome are currently known, while the remaining 85-90% of the estimated 600,000 PPIs remain to be discovered; for more than 5,000 of the human proteins, not even a single PPI is currently known (17). Discovery of PPIs is, therefore, essential to advancing research in biomedical science. Performing wet-lab experiments to detect *all* of these “missing” interactions is currently impossible due to limitations on funds, reagents, technical methods, and expertise over individual proteins. It becomes imperative that these unknown PPIs be predicted with computational methods or be detected with high-throughput biotechnological methods. We developed a computational model, called HIPPIP (high-precision protein-protein interaction prediction) that was deemed accurate by computational evaluations and experiments (18). Novel PPIs predicted using this model are making translational impact. For example, by constructing interactomes (i.e. PPI networks) of various disease associated genes, we have highlighted the role of cilia in congenital heart disease (19) and the role of mitochondrial proteins in hypoplastic left heart syndrome (20). Even individual PPIs have a potential to make vast impact. For example, the PPI between *oligoadenylate synthetase like* protein (OASL) and *retinoic acid-inducible gene I* (RIG-I) protein led to the discovery that OASL activates host response through RIG-I signaling during viral infections (21).

A number of databases and web applications make protein annotations and interactions available to biologists. Typically these web-based resources present comprehensive annotations of genes including their protein interactions (e.g., GeneCards, Wiki Genes, UniProt); some are exclusively designed for presenting lists of protein interactions of genes and also present network view of the PPIs (e.g., BioGRID, STRING,…). We developed a web resource called Wiki-Pi that presents individual PPIs with comprehensive annotations of the two proteins involved in an interaction (22). It allows search and retrieval of PPIs of interest based on their biological annotations and has helped several biologists interpret their experimental results. For example, it helped show a link between downregulation of the gene NDRG1 in oligodendrocytes of multiple sclerosis patients and altered translation of myelin genes resulting in partial demyelination (23) and pointing at the role of KPNA1 in regulating gene expression (24).

Our goal here is to apply HiPPIP to predict novel PPIs of MPM associated genes, and to make their predicted as well as previously known PPIs (namely, the MPM interactome) available in Wiki-Pi, so as to accelerate biomedical research on MPM. We demonstrate here the various ways in which system level analyses of the MPM interactome could lead to biologically and clinically relevant translatable results.

## Results

We collected MPM-associated genes from Ingenuity Pathway Analysis (IPA) suite, which gave a list of 62 genes curated from literature, which will be referred to as ‘MPM genes’ here; these genes have been reported to affect MPM through gene expression changes, or by harboring genetic variants, or by being targeted by drugs proven to be clinically active against MPM (see details in Data File 1) (25). PPIs of the 62 MPM genes were collected from two literature-curated databases, namely Human Protein Reference Database (HPRD) (26) and the Biological General Repository for Interaction Datasets (BioGRID) (27). Next, we applied HiPPIP, a computational model that we developed previously to predict novel PPIs of MPM genes. In HiPPIP, PPIs are predicted by computing features of protein pairs and developing a random forest model to classify the pairwise features as *interacting* or *non-interacting.* Protein annotations that were used in this work are: cellular localization, molecular function and biological process membership, genomic location of the gene, gene expression in hundreds of microarray experiments, protein domains and tissue membership of proteins. Computation of features of protein-pairs is described earlier in (28). A random forest with 30 trees was trained using the feature offering maximum information gain out of 4 random features to split each node; minimum number of samples in each leaf node was set to be 10. The random forest outputs a continuous valued score in the range of [0,1]. The threshold to assign a final label was varied over the range of the score for positive class (i.e., 0 to 1) to find the precision and recall combinations that are observed. This prediction model is referred to as High-precision Protein-Protein Interaction Prediction (HiPPIP) model. Applying HiPPIP, we discovered 367 novel PPIs of MPM genes, which are deemed highly accurate according to prior evaluation, including experimental validation, of the HiPPIP model (18). The MPM interactome has 1,387 known PPIs and 367 novel PPIs among the 62 MPM-associated genes and 1,620 interactors (Figure 1 and Data File 2). Nearly half of the MPM genes had fewer than 10 known PPIs each, whereas 150 novel PPIs are predicted for these together (Figure 2).

**Figure 1.**
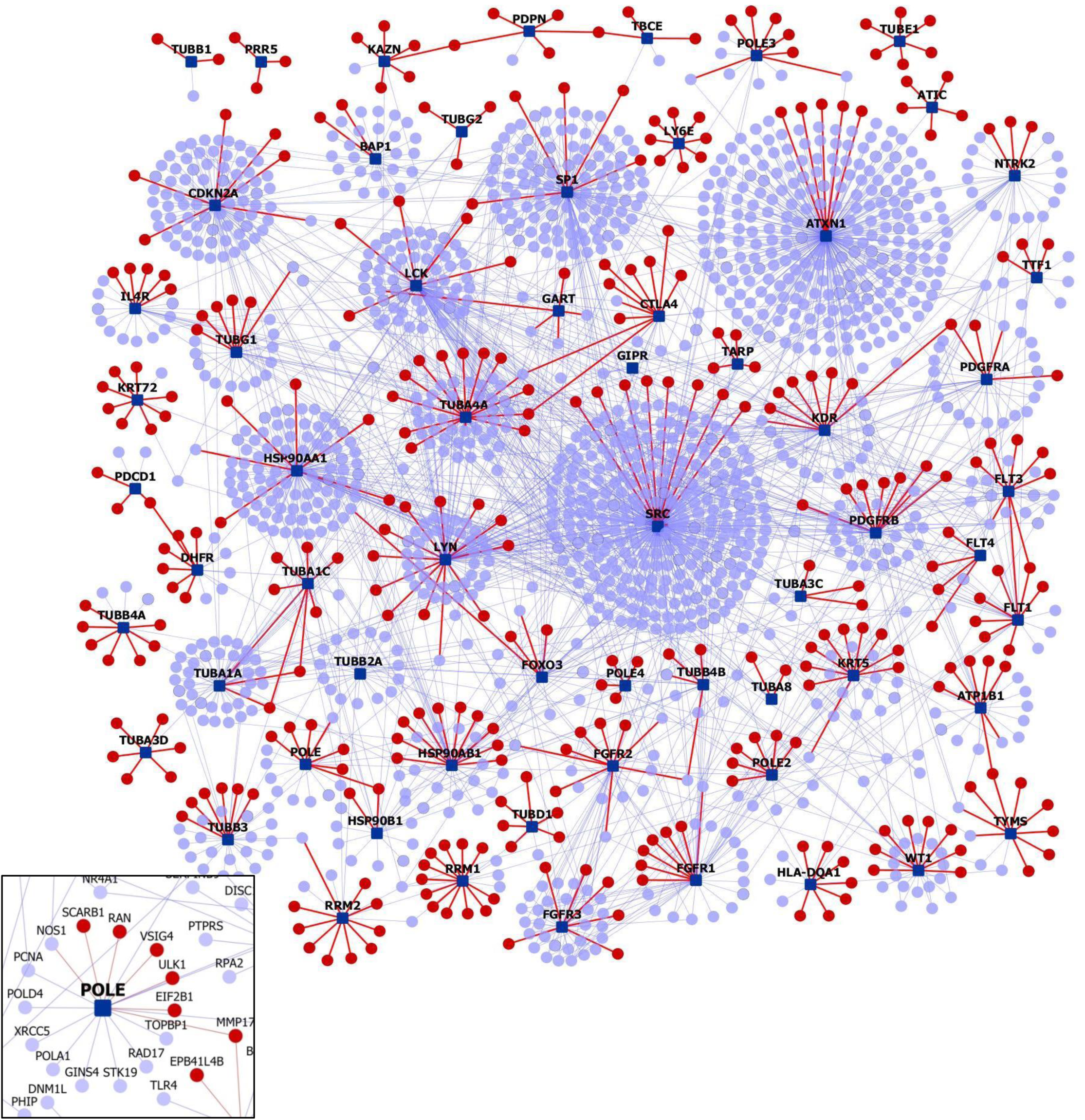
MPM Protein-Protein Interactome: Network view of the MPM interactome is shown as a graph where genes are shown as nodes and PPIs as edges connecting the nodes. MPM-associated genes are shown as dark blue square-shaped nodes, novel interactors and known interactors as red and light blue color circular nodes respectively. Red edges are the novel interactions, whereas blue edges are known interactions. The inset shows the connections in the

**Figure 2.**
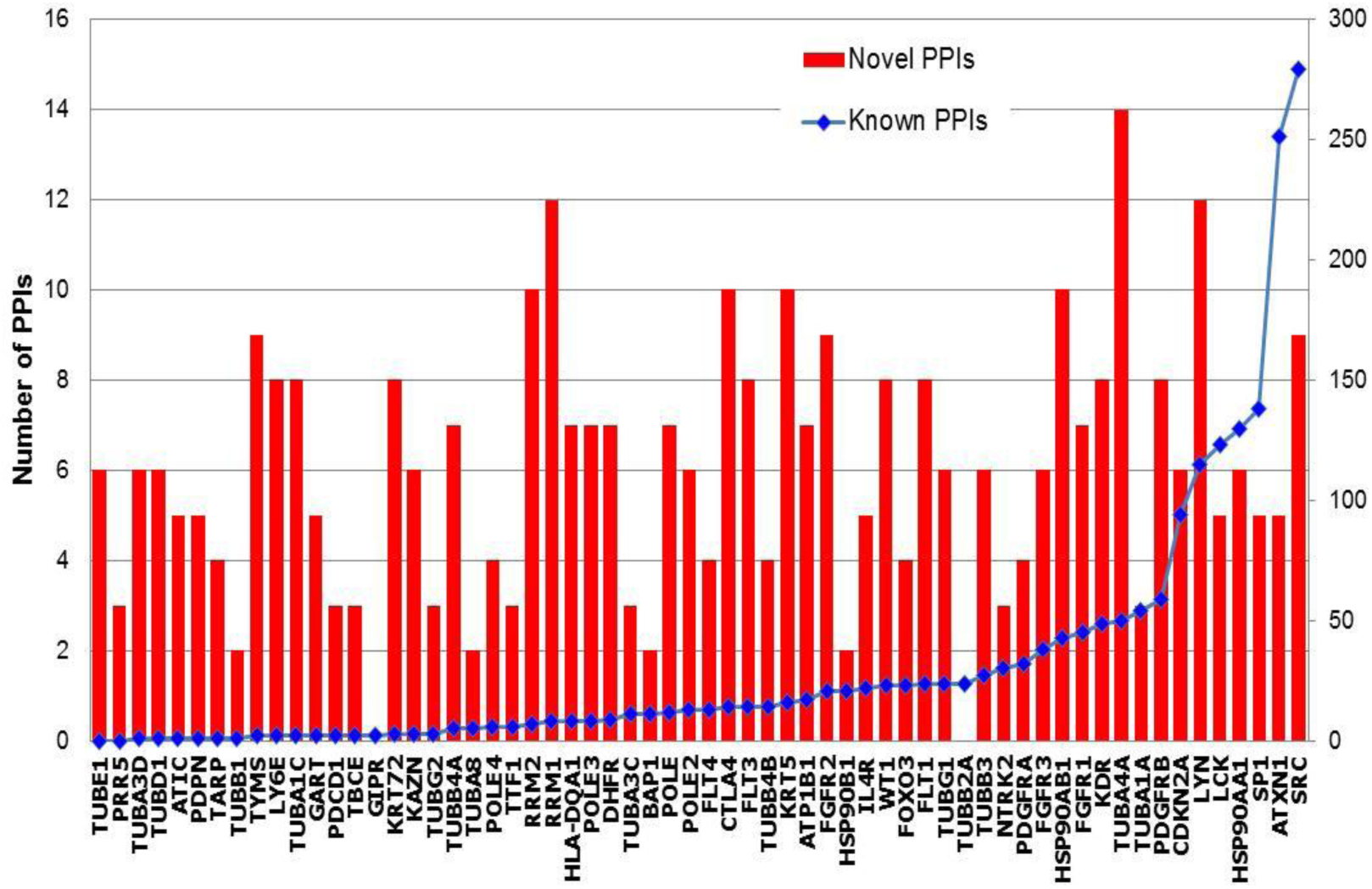
Number of PPIs in the MPM Interactome: The 62 MPM genes are shown along the X-axis, arranged in ascending order of their number of known PPIs. **Blue line, right-side axis:** Number of known PPIs is shown. **Red bars, left-side axis:** Number of novel PPIs.

### Experimental Validation of Selected PPIs

We carried out experimental validations of five predicted PPIs, namely, BAP1-PARP3, ALB-KDR, ALB-PDGFRA, CUTA-HMGB1 and CUTA-CLPS. These PPIs were chosen for their biological relevance as well as to their proximity to MPM genes. The first three are novel interactions of MPM genes, and the last two are in close proximity to multiple MPM genes in this biophysical interaction network. All 5 PPIs were validated using protein pull-down followed by protein identification. The pulled down sample is analyzed with either mass spectrometry (SF Table 1) or size-based protein detection assay (see Methods) to identify whether the prey protein has been pulled down along with the bait protein. Each bait protein was also paired with a random prey protein to serve as control (specifically, BAP1-phospholambin, ALB-FGFR2 and CUTA-FGFR2). All predicted PPIs but validated to be true, while control pairs tested negative. In addition to these 5 PPIs, another PPI from MPM interactome, namely HMGB1-FLT1 was validated in our prior work through co-immunoprecipitation (18).

**Table 1.** Pathways that are relevant to the pathophysiology and genetics of malignant pleural mesothelioma: Pathway analysis revealed that molecular mechanisms underlying various types of cancers, axonal guidance signaling, PI3/AKT signaling, VEGF signaling, natural killer cell signaling and inflammation signaling pathways may be pertinent to the development of MPM, among other pathways. A list of all the associated pathways and the genes from the interactome with each of these pathways is shown in Supplementary File 6.

| Pathway | P-value | Candidate genes | Novel genes |
| --- | --- | --- | --- |
| Glucocorticoid Receptor Signaling | 6.31E-56 | KRT72, HSP90B1, FGFR3, HSP90AB1, FGFR1, KRT5, FOXO3, FGFR2, HSP90AA1 | KRT74, HMGB1, PRKAG1, IL6, KRT6B, KRT78, KRT80, KRT7, KRT4, TAF3, NPPA, MAP3K7, KRT6A |
| Molecular Mechanisms of Cancer | 5.01E-53 | CDKN2A, SRC, FGFR3, FGFR1, FGFR2 | CDK14, CDK20, CDKN2B, PRKAG1, WNT7B, RND2, WNT6, CDK8, RHOQ, RAPGEF3, MAP3K7, WNT3 |
| NF-κB Signaling | 1.26E-39 | FGFR1, LCK, FLT1, KDR, PDGFRA, FGFR2, NTRK2, FGFR3, PDGFRB, FLT4 | MAP3K7 |
| Small Cell Lung Cancer Signaling | 2.00E-37 | FGFR1, FGFR2, FGFR3 | CDKN2B |
| Axonal Guidance Signaling | 2.51E-37 | TUBB1, TUBA1A, TUBA4A, TUBA8, TUBB2A, NTRK2, FGFR3, FGFR1, TUBB3, TUBG1, TUBA1C, TUBB4B, FGFR2, TUBB4A | MYL6B, DPYSL2, PRKAG1, PLCL1, WNT7B, WNT6, PLXNA1, TUBA3E, WNT3 |
| PI3K/AKT Signaling | 1.58E-36 | HSP90B1, FOXO3, HSP90AA1, HSP90AB1 | OCRL, PPP2CB, MCL1, EIF4EBP1 |
| VEGF Signaling | 3.98E-36 | FGFR1, FLT1, SRC, KDR, FOXO3, FGFR2, FGFR3, FLT4 | EIF2B1, EIF2B2 |
| Role of Macrophages, Fibroblasts and Endothelial Cells in Rheumatoid Arthritis | 6.31E-36 | SRC, FGFR3, FGFR1, FGFR2 | IL1RL1, IL6, PLCL1, WNT7B, IL7, WNT6, CALML5, MAP3K7, WNT3, APC2 |
| Natural Killer Cell Signaling | 6.31E-32 | FGFR1, LCK, FGFR2, FGFR3 | OCRL, CD244 |
| Actin Cytoskeleton Signaling | 1.58E-30 | FGFR1, FGFR2, FGFR3 | MYL6B, GSN, APC2 |
| eNOS Signaling | 3.16E-30 | FGFR1, FLT1, KDR, HSP90B1, FGFR2, HSP90AA1, FGFR3, FLT4, HSP90AB1 | PRKAG1, CALML5, AQP5, CHRM3 |
| Neuroinflammation Signaling Pathway | 3.98E-30 | FGFR1, HLA-DQA1, FGFR2, FGFR3 | HMGB1, HLA-DQB1, ACVR1B, IL6, GRIN1, GRIA1 |
| Gap Junction Signaling | 1.00E-29 | FGFR1, TUBB3, TUBG1, TUBB1, TUBA1C, TUBA1A, SRC, TUBB4B, TUBA4A, FGFR2, TUBA8, TUBB2A, FGFR3, SP1, TUBB4A | GJB3, PRKAG1, TUBA3E, PLCL1, GRIA1 |
| Integrin Signaling | 1.58E-28 | FGFR1, SRC, FGFR2, FGFR3 | GSN, ITGAM, RHOQ, CAPN2, RND2 |
| IL-6 Signaling | 1.58E-28 | FGFR1, FGFR2, FGFR3 | IL1RL1, MCL1, IL6, MAP3K7 |

We hypothesize that the BAP1-PARP3 interaction may enhance cancer growth in MPM. BAP1 is a tumor suppressor protein playing a role in cell cycle progression, repair of DNA breaks, chromatin remodeling, and gene expression regulation, and variants in BAP1 have been implicated in hereditary and sporadic mesothelioma (29). PARP3 is involved in DNA repair, regulation of apoptosis and maintenance of genomic stability by stabilizing the mitotic spindle, and maintaining telomere integrity (30). In clinical trials (31), PARP inhibitors were found to influence cancers in which mutations in BRCA1 or BRCA2 are observed. BAP1 interacts with BRCA1, another tumor suppressor protein, inhibiting breast cancer growth (14).(14)(14)(14)(14) In the absence of BRCA1 activity or with a perturbation in its interaction with BAP1, cancerous growth is enhanced (31). Then, the novel interaction of BAP1 with PARP3 in cancerous cells may promote cancerous growth, possibly through regulation of DNA repair and apoptosis. BAP1and PARP3 were found to be mildly over expressed in sarcomatoid MPM tumors compared with normal pleural tissue (log2(fold change) or log2FC=0.575, p-value=0.028, log2FC=0.695, p-value=0.0212, respectively) (GSE42977 (32)). Low levels of ALB have been correlated with poor prognosis in MPM patients (33). The two MPM genes, KDR and PDGFRA, that ALB is predicted to interact with, are members of the PI3K/AKT pathway which is aberrantly active in mesothelioma (34). High expression of CUTA has been correlated with favorable prognosis in lung cancer (Pathology Atlas). It was found to be overexpressed in MPM tumors vs. normal pleura (log2FC = 0.871, p-value = 0.0039) (GSE2549 (35)) and in MPM tumors vs. other thoracic tumors (log2FC = 0.454, p-value = 0.0029) (GSE42977 (32)). CLPS inhibits metastasis of the melanoma cell line, B16F10, to lungs by blocking the signaling pathway involving β1 integrin, FAK and paxillin (36). CLPS has a novel interaction with NEDD9, which has been shown to mediate β1 integrin signaling and promote metastasis of non-small lung cancer cells(37). CD26, a cancer stem cell marker of malignant mesothelioma, has been shown to associate with the integrin α5β1 (or ITGA5, a novel interactor of the MPM gene, FGFR2) integrin and promote cell migration and invasion in mesothelioma cells (37). Another cancer stem cell marker of malignant mesothelioma, CD9, inhibits this metastatic effect mediated by CD26. Depletion of CD26 and CD9 was shown to lead to decreased and increased expression of NEDD9 and FAK in mesothelioma cells lines, hinting at the involvement of NEDD9 in mesothelioma tumor invasiveness (37). NEDD9 has a known interaction with LYN, an MPM gene, shown to play a negative role in the regulation of integrin signaling in neutrophils (38). CUTA has a novel interaction with HMGB1, which has been shown to activate the integrin αMβ2 (or ITGAM, a novel interactor of the MPM gene, TYMS) and the cell adhesion and migratory function of neutrophils mediated by αMβ2 (39). HMGB1 also has a novel interaction with the MPM gene, FLT1, shown to be involved in the migration of multiple myeloma cells by associating with β1 integrin, and mediating PKC activation (40).

### Web Server

We made the MPM interactome available on a webserver called *Wiki-Pi MPM* (http://severus.dbmi.pitt.edu/wiki-MPM). It has advanced-search capabilities, and presents comprehensive annotations, namely their Gene Ontology annotations, diseases, drugs and pathways, of the two proteins of each PPI side-by-side. Here, a user can query for results such as “show me PPIs where one protein is involved in mesothelioma and the other is involved in immunity”, and then see the results with the functional details of the two proteins side-by-side. The PPIs and their annotations also get indexed in major search engines like Google and Bing, thus a user searching for ‘KDR and response to starvation’ would find the PPIs KDR-CAV1 and KDR-ALB, where the interactors are each involved in ‘response to starvation’. This is a unique feature of this web server, as such results are not supported by any other PPI web database. The novel PPIs have a potential to accelerate biomedical discovery in mesothelioma, and making them available on this web server brings them to the biologists in an easily-discoverable and usable manner.

### Analysis with other High-Throughput Data

The interactome of MPM genes may serve as a vehicle to piece together an integrated view of functional interconnections among genes linked to MPM through various high throughput studies. We analyzed the overlap of the interactome with various high throughput datasets such as MPM-associated genetic variants, differential expression, and methylation in MPM, correlation of gene expression with lung cancer prognosis and differential gene expression on asbestos exposure, details of which are presented here.

We collected the genetic variants identified through mutational profiling of MPM tumors in Bueno et al. (41) and found that 253 genes in the MPM interactome had either germline mutations, or somatic single nucleotide variants (SNVs) or indels (insertions or deletions) (Data File 3). Of these 253 genes, 39 were novel interactors of MPM genes: MGMT carried germline mutations while the following carried somatic mutations: KIAA1524, SLC20A1, LATS2, ASTN2, BARX1, BRD2, CALML5, CAPRIN1, CLK1, CPS1, DPYD, EIF3H, EPB41L3, GMPS, GPR12, ITGAM, KMT2D, KRT4, MGAT4A, NBR2, NDUFV2, NFIB, NFX1, NUDC, PLCL1, PRKAG1, PRMT1, PTPRT, PTRH2, RBBP6, SGK3, SMCHD1, SPOCK1, TMPRSS15, TNC, XPO4, ZNF687 and PRDM2 carried somatic mutations. Of these 39 novel interactors, twelve interact with MPM genes that also harbored a genetic variant. The PPIs in which both genes carried genetic variants are: CDKN2A-NFX1, FLT1-LATS2, TUBA3C-XPO4, PDGFRA-SPOCK1, TYMS-SMCHD1, TYMS-EPB41L3, GART-TMPRSS15, TYMS-NDUFV2, TYMS-ITGAM, RRM2-BARX1, RRM2-MGAT4A and ATIC-CPS1.

We collected the methylation profile of pleural mesothelioma (42), and found nine novel interactors to be hypomethylated in pleural mesothelioma compared with non-tumor pleural tissue, namely, ACVR1B, CDKN2B, IL6, MGMT, NRG1, OAT, PHLDA2, PLAUR and TNC (SF Table 2). Some of them have little or no expression in lung tissue but are overexpressed in MPM. PLAUR is a prognostic biomarker of human MPM, and is also associated with shorter survival time in a rat xenograft model of malignant mesothelioma (43). Similarly, FGFR1 and its novel interactor NRG1 had elevated mRNA expression in H2722 mesothelioma cell lines and in MPM tissue, both contributing to increased cell growth under tumorigenic conditions (44, 45). TNC, which contributes to invasive growth, is a prognostic biomarker overexpressed in MPM and in malignant pleural effusion, having low expression in normal lung tissues(46, 47). Thus, these novel interactors, which are not normally expressed in lung tissue, may be hypomethylated in MPM leading to their over-expression, contributing to MPM etiology. We computed the overlap of genes differentially expressed in mesothelioma tumors vs. normal pleural tissue adjacent to tumor (GSE12345 (48)) and found statistically significant overlap with the MPM interactome (genes with fold-change >2 or <½ was considered as overexpressed and under expressed respectively at a P-value<0.05). 336 genes out of the 1,682 genes in the MPM interactome overlapped with this dataset (p-value of overlap=2.263e-13), out of which 53 genes were novel interactors. Similarly, 656 out of the 1,682 genes in the MPM interactome were differentially expressed in MPM tumors vs. other thoracic cancers such as thymoma and thyroid cancer (GSE42977 (32) (p-value of overlap = 1.36E-42). 113 out of the 367 novel interactors were differentially expressed in this dataset (p-value of overlap = 0.0034). This shows that the MPM interactome is enriched with genes whose expression helps in distinguishing MPM from other thoracic tumors and also with genes differentially expressed in mesothelioma tumors compared with adjacent normal pleural tissue (Figure 3 and Data File 3). 325 genes in the MPM interactome have high/medium expression in normal lung tissue (median transcripts-per-million (TPM)≥9), using RNA-sequencing data available in GTEx (Figure 3 and Data File 3) (49). Of these, 61 were novel interactors.

**Table 2.** Pharmacokinetic details of known mesothelioma drugs and the drugs that are presented as candidates for repurposing and their correlation scores with lung cancer expression studies from NextBio analysis: Pharmacokinetic details are from Drug Bank (https://www.drugbank.ca/). Known mesothelioma drugs are shown in italics. The correlation score is based on the strength of the overlap or enrichment between the two biosets. Additional statistical criteria such as correction for multiple hypothesis testing are applied and the correlated biosets are then ranked by statistical significance. A numerical score of 100 is assigned to the most significant result, and the scores of the other results are normalized with respect to the top-ranked result.

| Drug name | Original therapeutic purpose(s) | Dosage form | Delivery route | Half-life | Toxicity | MPM Targets | Correlation of lung cancer expression with drug | Correlation score |
| --- | --- | --- | --- | --- | --- | --- | --- | --- |
| *Pemetrexed* | Chemotherapeutic drug for pleural mesothelioma and non-small cell lung cancer | Powder for solution | Intravenous | 3.5 hours | Data not available | ATIC, DHFR, GART, TYMS | negative | 79 |
| *Mitomycin* | Chemotherapeutic drug for breast, bladder, oesophageal, stomach, pancreas, lung and liver cancers | Injection, powder or lyophilized for solution | Intravenous | 8-48 minutes | Nausea and vomiting | - | negative | 64 |
| Cabazitaxel | Anti-neoplastic agent in hormone-refractory metastatic prostate cancer | Solution | Intravenous | Rapid initial-phase of 4 mins, intermediate-phase of 2 hours and prolonged terminal-phase of 95 hours | Neutropenia, hypersensitivity reactions, gastrointestinal symptoms, renal failure | TUBB1, TUBA4A | negative | 79 |
| Pyrimethamine | Anti-parasitic agent in toxoplasmosis and acute malaria | Tablet | Oral | 4 days | Data not available | DHFR | negative | 83 |
| Trimethoprim | Anti-bacterial agent/antibiotic in urinary tract, respiratory tract and middle-ear infections and traveller's diarrhea | Tablet/solution | Oral | 8 to 11 hours | Oral toxicity in mice at LD50=4850 mg/kg | DHFR, TYMS | negative | 63 |
| Primaquine | Anti-malarial agent | Tablet | Oral | 3.7 to 7.4 hours | Data not available | KRT7 | negative | 71 |
| Gliclazide | Anti-diabetic/hypoglycemic medication in type 2 diabetes mellitus | Tablet | Oral | 10.4 hours | Oral toxicity in mice at LD50=3000 mg/kg, accumulation in people with severe hepatic and/or renal dysfunction, side-effects of hypoglycemia including dizziness, lack of energy, drowsiness, headache and sweating | VEGFA | negative | 56 |

**Figure 3.**
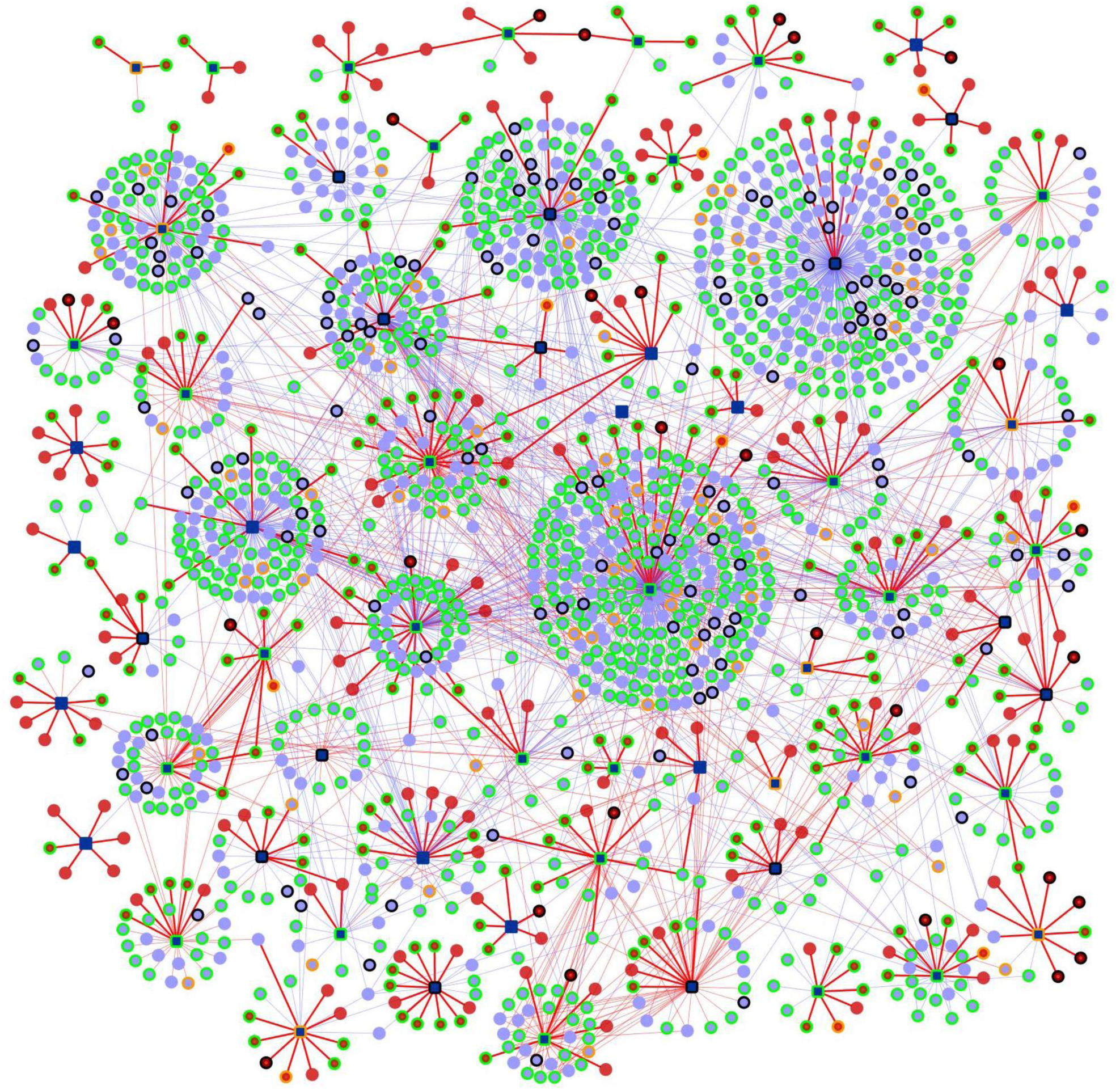
Genes with biological evidences in the MPM Protein-Protein Interactome: On the interactome network shown in Figure 1, various biological evidences of relation to MPM are shown as node border colors. Genes with variants associated with MPM have orange borders, genes with MPM/lung cancer/asbestos exposure-associated gene expression changes have light green-colored borders and genes with black border have both genetic variants and gene expression changes; the features include differential expression in MPM tumors vs. normal adjacent pleura, MPM tumors vs. other thoracic tumors, differential gene methylation (affecting gene expression) in MPM tumors, gene expression correlated with unfavorable lung cancer prognosis and differential gene expression on exposure to asbestos or asbestos-like particles

According to Pathology Atlas data, 63 genes from the interactome (of which are 10 novel interactors) were found to be those whose high expression was positively correlated with unfavorable prognosis for lung cancer, namely, SPOCK1, SLC7A5, SCARB1, PLIN3, PLAUR, PIEZO1, KRT6A, GJB3, B3GNT3 and ARL2BP (2.08 odds ratio; p-value =1.91E-08) (50). In small cell lung cancer cells, expression of ARL2BP has been shown to be associated with high expression of YAP, which has oncogenic action in lung cancer (Data File 2)(51). In our interactome, ARL2BP has been predicted to interact with FLT1, a VEGF receptor expressed in MPM cells. VEGF levels in the serum of MPM patients has been inversely correlated with patient survival, i.e. unfavorable prognosis, and non-small cell lung cancer tumors expressing FLT1 have been associated with higher malignancy and poor prognosis (52, 53).

A recent study has reported the genes that were differentially expressed in lungs of mice exposed to crocidolite and erionite fibers compared to a control group (54). Crocidolite and erionite are asbestos or asbestos-like particles that are capable of inducing lung cancer and/or mesothelioma in humans and animal models (54). Out of the 1,710 genes differentially expressed on crocidolite exposure, 160 genes were part of the MPM interactome (p-value of overlap =3.63E-05), and 6 out of 78 genes differentially expressed on erionite exposure occured in the MPM interactome (Figure 4 and Data File 3).

**Figure 4.**
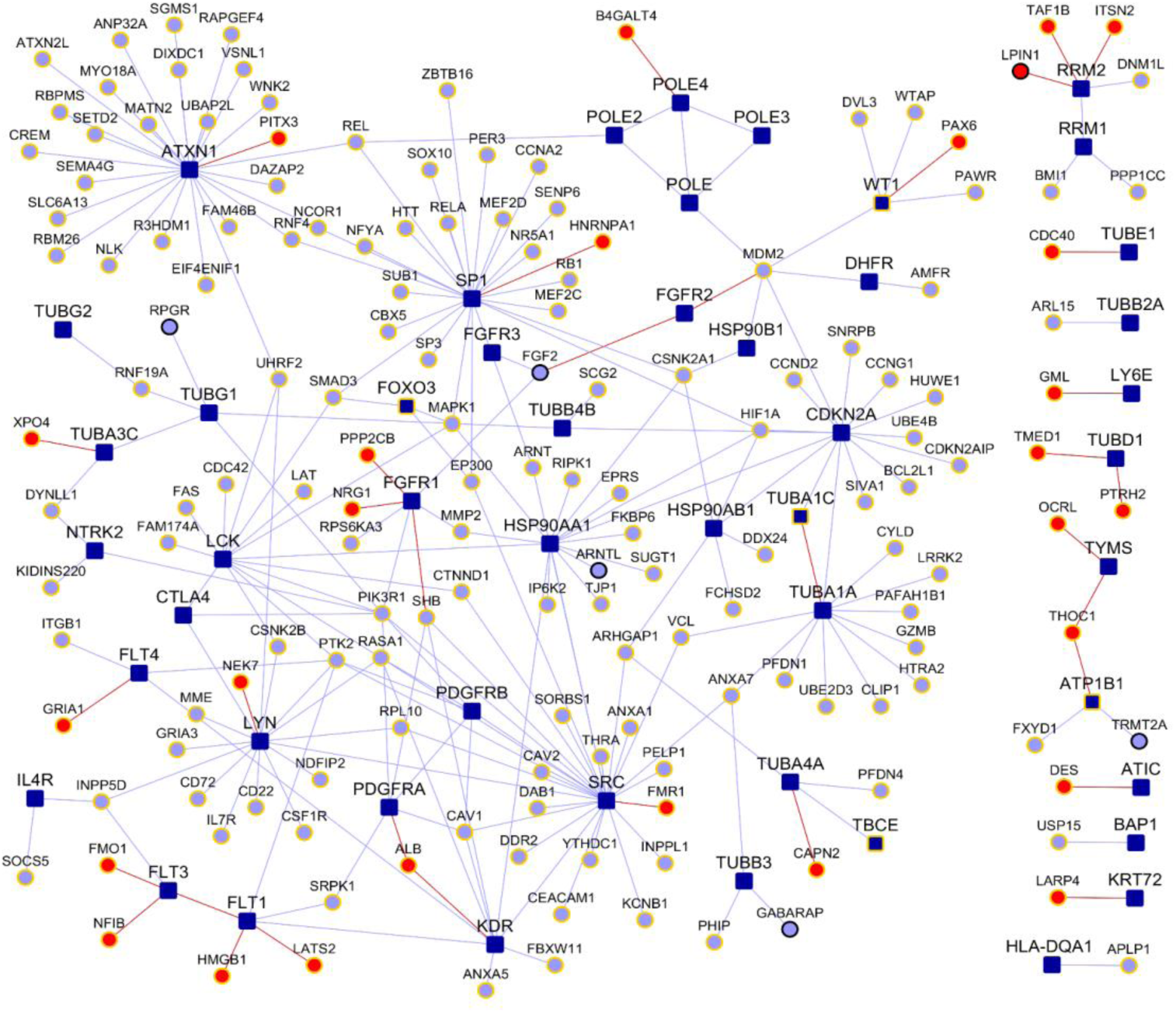
MPM Interactome genes that are differentially expressed upon particle exposure: Gene fill colors and edge colors are the same as in the main interactome (Figure 1), whereas border colors indicate differential expression in exposure to crocidolite (yellow border) and erionite (black border).

### Pathway analysis

We compiled the list of pathways that any of the proteins of MPM interactome are associated with, using IPA suite (55). A number of pathways having highly significant association with genes in the MPM interactome such as *NF-κB* Signaling (p-value=1.25E-39), *PI3/AKT signaling* (p-value= 1.58E-36), *VEGF signaling* (p-value=3.98E-36) and *natural killer cell signaling* (p-value=6.3E-32) were identified (Table 1 and Data File 4). Top 30 pathways by statistical significance of association are also shown in Figure 5A. These pathways are highly relevant to mesothelioma etiology. For example, the PI3K/AKT signaling pathway, which regulates the cell cycle and is involved in cell proliferation, becomes aberrantly active in MPM (Figure 5B) (56). The supplementary data (Data Files 2 and 4) made available here allows a cancer biologist to study PPIs, including hitherto unknown novel PPIs, that connect MPM genes to a pathway that they are interested in studying.

**Figure 5.**
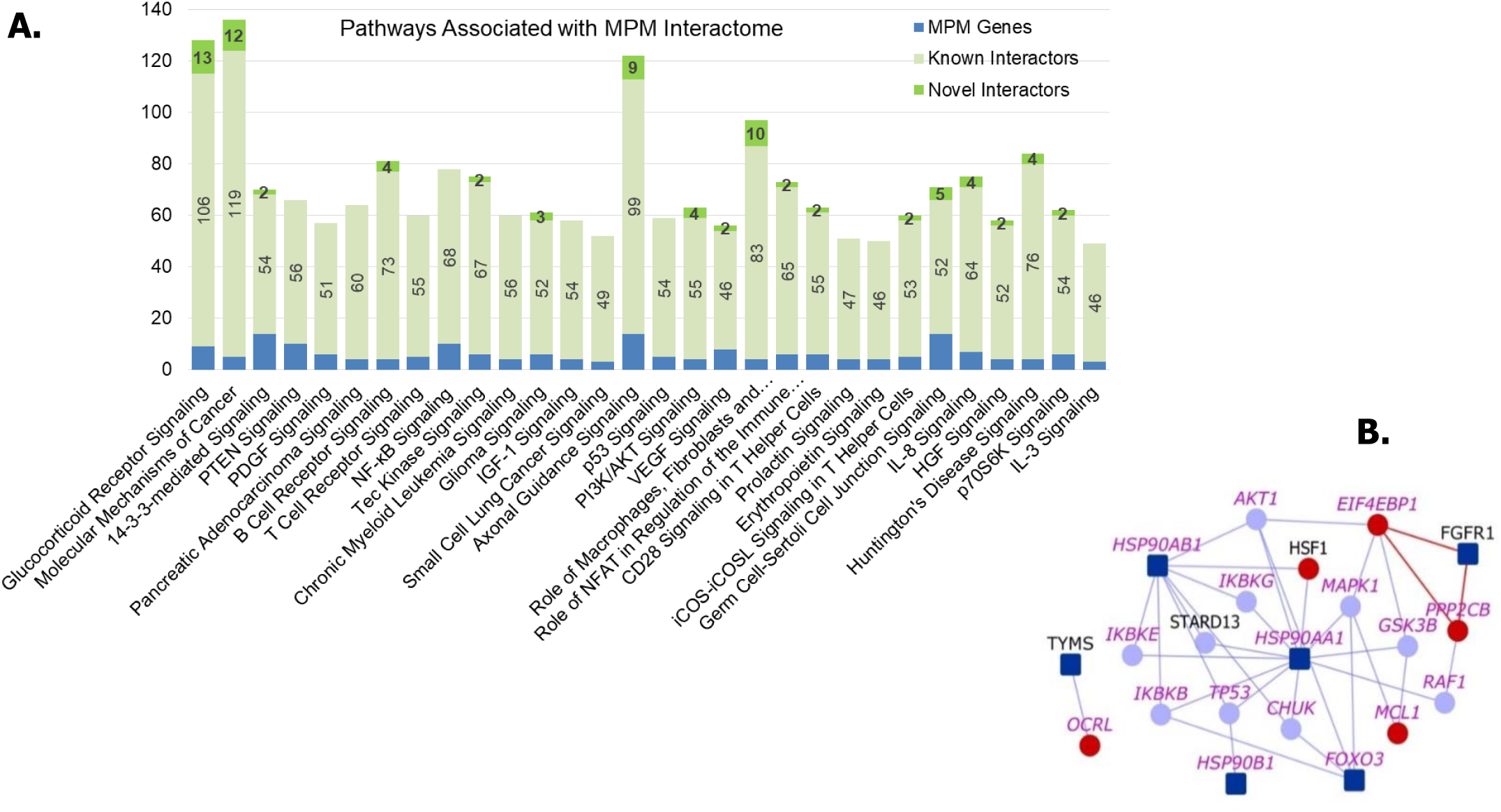
Pathways associated with MPM interactome: (A) Number of genes from MPM interactome associated with various pathways are shown. Top 30 pathways based on significance of association with the interactome are shown. (B) PI3K/AKT Signaling Pathway: Dark blue nodes are MPM genes, light blue nodes are known interactors and red nodes are novel interactors. Nodes with purple labels are genes involved in the PI3K/AKT signaling pathway.

### Potentially Repurposable Drugs

Our previous work on schizophrenia interactome analysis led to the identification of drugs potentially repurposable for schizophrenia of which one of them is currently in clinical trials (57). Following this methodology, we constructed the MPM drug-protein interactome that shows the drugs that target any protein in the MPM interactome. There are 513 unique drugs that target 206 of these proteins (of which 28 are novel interactors that are targeted by 147 drugs) (Figure 6 and Data File 5). Using an approach of comparing differential expression induced by the disease versus a drug (58), we identified five drugs that could be potentially repurposable for MPM. These are: *cabazitaxel*, a cancer drug used in the treatment of refractory prostate cancer; *primaquine* and *pyrimethamine*, two anti-parasitic drugs; *trimethoprim*, an antibiotic; and *gliclazide*, an anti-diabetic drug. Method used for identifying repurposable drugs has been detailed in Supplementary Methods.

**Figure 6.**
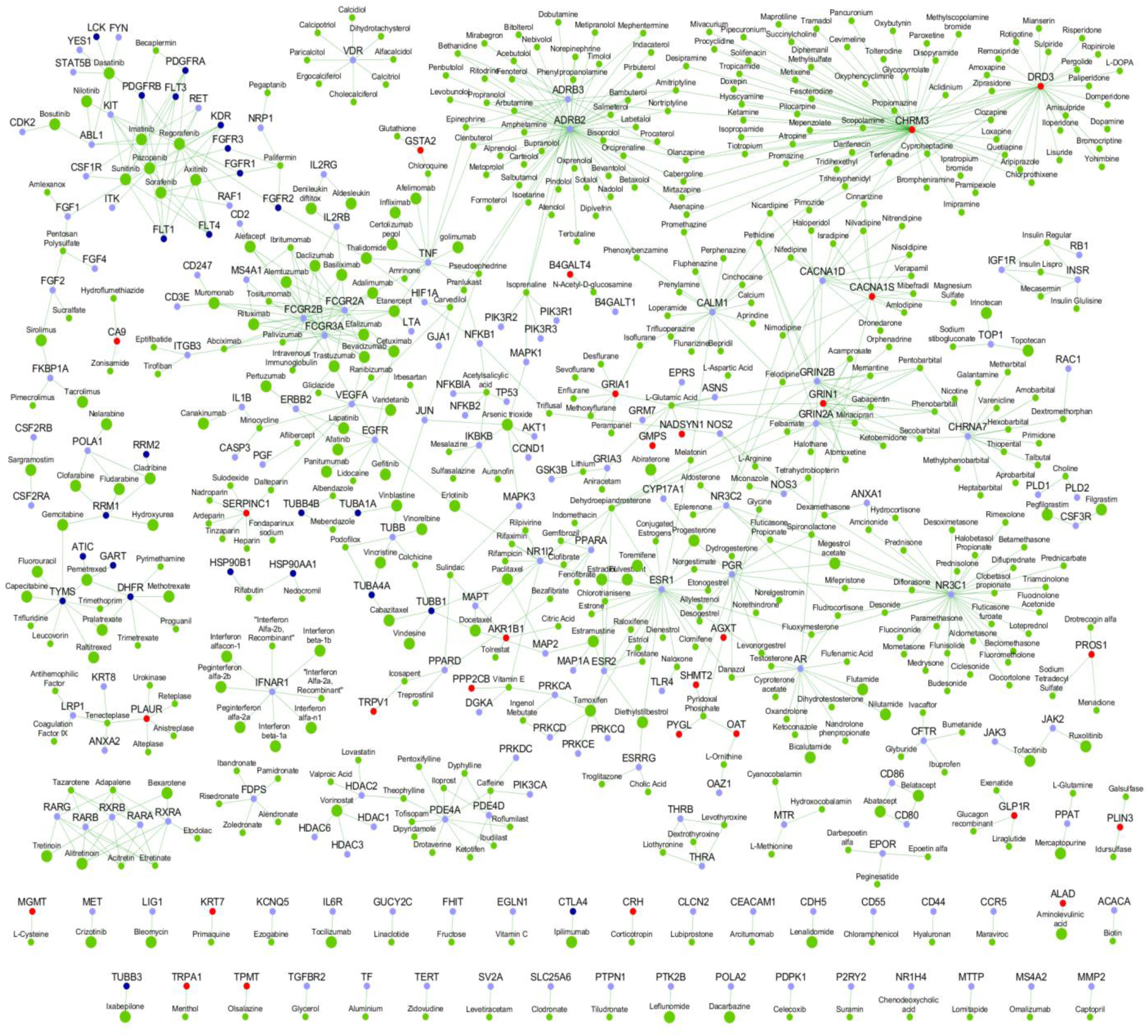
MPM Drug-Protein Interactome: The network shows the drugs (green color nodes) that target the proteins in the MPM interactome. Larger green nodes correspond to drugs that target the anatomic category ‘antineoplastic and immunomodulating agents’ The color legend for genes (proteins) is as shown as in Figures 1, with MPM genes in dark blue, their known interactors in light blue and novel interactors in red.

We adopted an approach of comparing differential expression induced by the disease versus a drug (58) using the BaseSpace correlation software (https://www.nextbio.com) (59) to identify five drugs that are potentially repurposable for MPM (Supplementary Methods). Drugs were selected based on whether they were already tested against lung cancer in clinical trials and/or showed overall negative correlation with lung cancer expression studies, because both mesothelioma and lung cancers have been shown to share common pathways that are initiated on exposure to asbestos fibres in mesothelial cells and lung epithelial cells respectively (1).

Another criterion used was whether the genes targeted by the drugs showed high differential expression in MPM tumours/cell lines (GSE51024 (*60)* and GSE2549 (35)). Although in each case, there would be some genes that are differentially expressed in the same direction for both the drug and disorder (i.e. both the drug & disease cause some genes to over-express; or both the drug and disease cause other genes to under express), the overall effect on the entire transcriptome has an anti-correlation. A correlation score is generated by the tool based on the strength of the overlap between the two datasets. Other statistical criteria such as correction for multiple hypothesis testing are applied and the correlated datasets are then ranked by statistical significance. A numerical score of 100 is assigned to the most significant result, and the scores of the other results are normalized with respect to the top-ranked result. We excluded drugs with unacceptable toxicity (e.g. minocycline) or unsuitable pharmacokinetics. The final list comprised of 15 drugs, out of which 11 have already been tested against mesothelioma in clinical trials/animal models and several of them were found to display clinical activity. Gemcitabine and pemetrexed are being used as first-line therapy for mesothelioma, in combination with cisplatin (61) (62). Ipilimumab has been identified to be a potential second- or third-line therapy in combination with nivolumab (63). Ixabepilone stabilizes cancer progression for up to 28 months (64) Zoledronate, which showed modest activity in MPM, induced apoptosis and S-phase arrest in human mesothelioma cells and inhibited tumor growth in the pleural cavity of an orthotopic animal model (65, 66). Sirolimus/Cisplatin increased cell death and decreased cell proliferation in cell lines of malignant pleural mesothelioma (MPM) (67). α-tocopheryl succinate increased survival of orthotopic animal models of malignant peritoneal mesothelioma (68). Testing of Vitamin E and its analogs are being carried out in various pre-clinical settings (69, 70). Eliminating those drugs which are being/have already been tested in mesothelioma with varying results, we arrived at a list of 5 potentially repurposable drugs in the descending order of negative correlation scores: pyrimethamine, cabazitaxel, primaquine, trimethoprim and gliclazide (Table 2). Cabazitaxel targets the MPM genes, TUBB1 and TUBA4A, and was effective in treating NSCLC that was resistant to docetaxel, a drug that targets TUBB1 along with other known interactors of MPM genes (71). Pyrimethamine and Trimethoprim target two MPM genes, involved in folate metabolism, that were highly differentially expressed in MPM tumors (GSE51024 (60)): TYMS (log2FC = 1.82, P-value = 4.10E-17) and DHFR (log2FC = 0.89, P-value =1.20E-14). MPM tumors have been shown to be responsive to anti-folates (72). Primaquine targets KRT7, a novel interactor of KRT5, whose high expression has been correlated with tumour aggressiveness and drug resistance in malignant mesothelioma (73-75). Primaquine may be re-purposed for MPM treatment at least as an adjunctive drug with pemetrexed, the drug currently used for first line therapy. Primaquine enhanced the sensitivity of the multi-drug resistant cell line KBV20C to cancer drugs (76). Gliclazide is an anti-diabetic drug inhibits VEGFA, a known interactor of KDR^8^, and is significantly upregulated in MPM tumour (Log2FC = 1.83, P-value = 0.0018). Glicazide inhibits neovascularization, a process mediated by VEGF (77). High levels of VEGF have been correlated with both asbestos exposure in MPM and an advanced stage of the disease (78, 79). Glibenclamide, a drug with a similar mechanism of action as that of glicazide, increases caspase activity in MPM cell lines and primary cultures, leading to apoptosis mediated by TRAIL (TNF-related apoptosis inducing ligand) (80).

## Discussion

Currently, only a handful of genes, such as BAP1, CDKN2A and NF2, are being actively studied for their relation to mesothelioma. In an effort to shed light onto other genes associated with MPM, whose functions are uncharacterized, we assembled the ‘MPM interactome’ with ~1,300 previously known PPIs and 367 computationally predicted PPIs. The primary objectives of this paper are to make the interactome and its annotations available to researchers and to demonstrate the power of interactome-scale analyses to generate biological results that contribute to understanding the underlying etiology as well as those that may be directly translated to clinical research. We demonstrated the biological validity of the interactome by showing that it had highly significant overlaps with relevant biological datasets such as MPM-associated genetic variants, differential expression and methylation in MPM, correlation of gene expression with lung cancer prognosis and differential gene expression on asbestos exposure. Next, pathway analysis on the interactome revealed several cancer-related pathways significantly enriched in the interactome. We then expanded the MPM interactome to include drugs that target the proteins in the interactome. An integrated computational analysis with this drug-protein interactome, based on comparing differential expression induced by the disease versus a drug led us to shortlist five potentially repurposable drugs for MPM - an example of a clinically translatable result.

The interactome accelerates discovery by revealing genes or pathways which are not intuitively associated with MPM but are valuable to understanding its underlying biology, such as the association of *axon guidance signaling* (p-value=2.51E-37), and by allowing biologists to formulate novel biological hypotheses. An illustrative example demonstrating the use of interactome and its annotations to formulate a novel hypothesis is shown in Figure 7. The hypothesis drawn here is that the interplay of genes in semaphorin signaling could be contributing to the development of MPM. While interactome level analysis revealed that axon signaling pathway is significantly associated with MPM, a closer look at an individual novel PPI (TUBA4A-DPYSL2) in this pathway helped us to *generate a testable hypothesis that inactivation of SEMA3A-mediated signaling may promote microtubule assembly and increase cell proliferation in MPM.*

**Figure 7.**
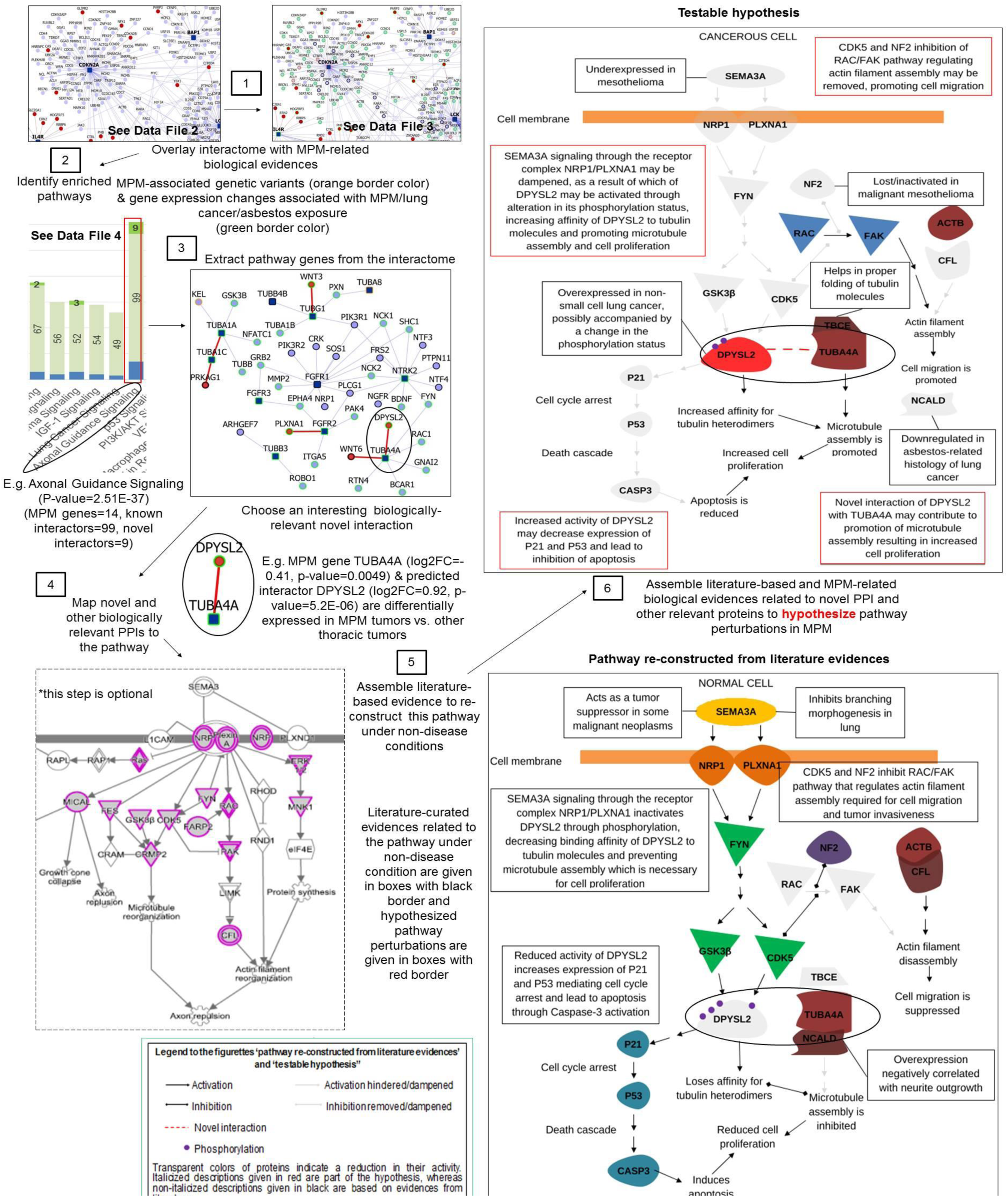
Illustrative example of how to utilize the MPM interactome and its annotations presented here, to formulate biological hypothesis. Evidences necessary for formulation of hypotheses are available in data files 2, 3 and 4, having predicted and known PPIs in the MPM interactome, MPM-related biological evidences corresponding to genes in the MPM interactome and pathways enriched in the interactome, respectively. Gene Ontology annotations, diseases, drugs and pathways of genes in the MPM interactome are available in Wiki-Pi MPM (http://severus.dbmi.pitt.edu/wiki-MPM).

Specifically, genes involved in axon guidance signaling were extracted from the interactome. Literature-based evidences were then assembled to re-construct these pathway-relevant interactions under non-disease conditions. Further MPM-related biological evidences (MPM-associated genetic variants and gene expression changes associated with MPM/lung cancer/asbestos exposure) related to novel PPI and other relevant proteins were considered from Figure 3 and Data File 2 to hypothesize pathway perturbations in MPM. Below, we present some testable hypotheses of novel interactions in the MPM interactome. Literature based study showed that semaphorins have been known to be involved in the development of non-neural organs and are also implicated in the repulsion of growth-cones of nerves during axon guidance (81). SEMA3A is an axon guidance molecule inhibiting branching morphogenesis in lung organ culture, expressed in airway and alveolar epithelial cells of lungs (82). It is a tumor suppressor gene, underexpressed in malignant mesothelioma and in non-small cell lung cancer (83, 84). DPYSL2 (‘CRMP2’) is a known mediator of SEMA3A, expressed during the embryonic and alveolar stages of lung development in mice, alluding to its role in pulmonary innervation and alveolarization (85). SEMA3A signaling through the receptor complex NRP1/PLXNA1 inactivates DPYSL2 by phosphorylation, decreasing its binding affinity to tubulin molecules, preventing microtubule assembly and thereby interfering with proliferation of cells (86). Conversely, lower expression of SEMA3A in malignant mesothelioma may allow activation of DPYSL2, resulting in increased proliferation. In line with this, DPYSL2 is overexpressed in mesothelioma and in non-small cell lung cancer, possibly accompanied by a change in the phosphorylation status (87). Its novel interaction with TUBA4A (Figure 8) may contribute to promotion of microtubule assembly and increased cell proliferation. Known interaction of TUBA4A with TBCE may contribute to microtubule assembly because TBCE has been shown to help in proper folding of β-tubulin molecules to maintain networks of neuronal microtubules (88). Overexpression of NCALD, a known interactor of TUBA4A, has been negatively correlated with neurite outgrowth (89). This interaction may be disrupted in MPM, as NCALD was found to be significantly down regulated in asbestos-related histology of lung cancer (90). Reduced activity of DPYSL2 also increase the expression of CDKNA1 (‘p21’) and TP53 (‘p53’) causing cell cycle arrest and apoptosis through activation of CASP3. Hence, increased DPYSL2 activity may inhibit apoptosis (87). CDK5, a downstream effector in SEMA3A-mediated signaling, and NF2, a tumor suppressor gene inactivated in MPM, inhibit the RAC/FAK pathway regulating actin filament assembly required for cell migration and tumor invasiveness (91). This inhibition on actin assembly may be removed due to dampening of SEMA3A-mediated signaling, leading to inactivation of CDK5 and independent inactivation of NF2 in MPM, promoting cell migration. Upregulation of the RAC/FAK pathway and increased tumor invasiveness have been linked to inactivation of NF2 in malignant mesothelioma (92). The interaction of ACTB which is usually synthesized in response to guidance cues, with CFL1 that promotes disassembly of actin filaments may also be disrupted in MPM (93, 94). Overexpression of CFL1 suppresses the growth of non-small cell lung cancer tumor, through suppression of cell motility and invasion (95). Thus, we conclude that the novel interaction of DPYSL2 with TUBA4A and its involvement in SEMA3A-mediated signaling may throw light on the role played by axon guidance signaling in mesothelioma.

**Figure 8.**
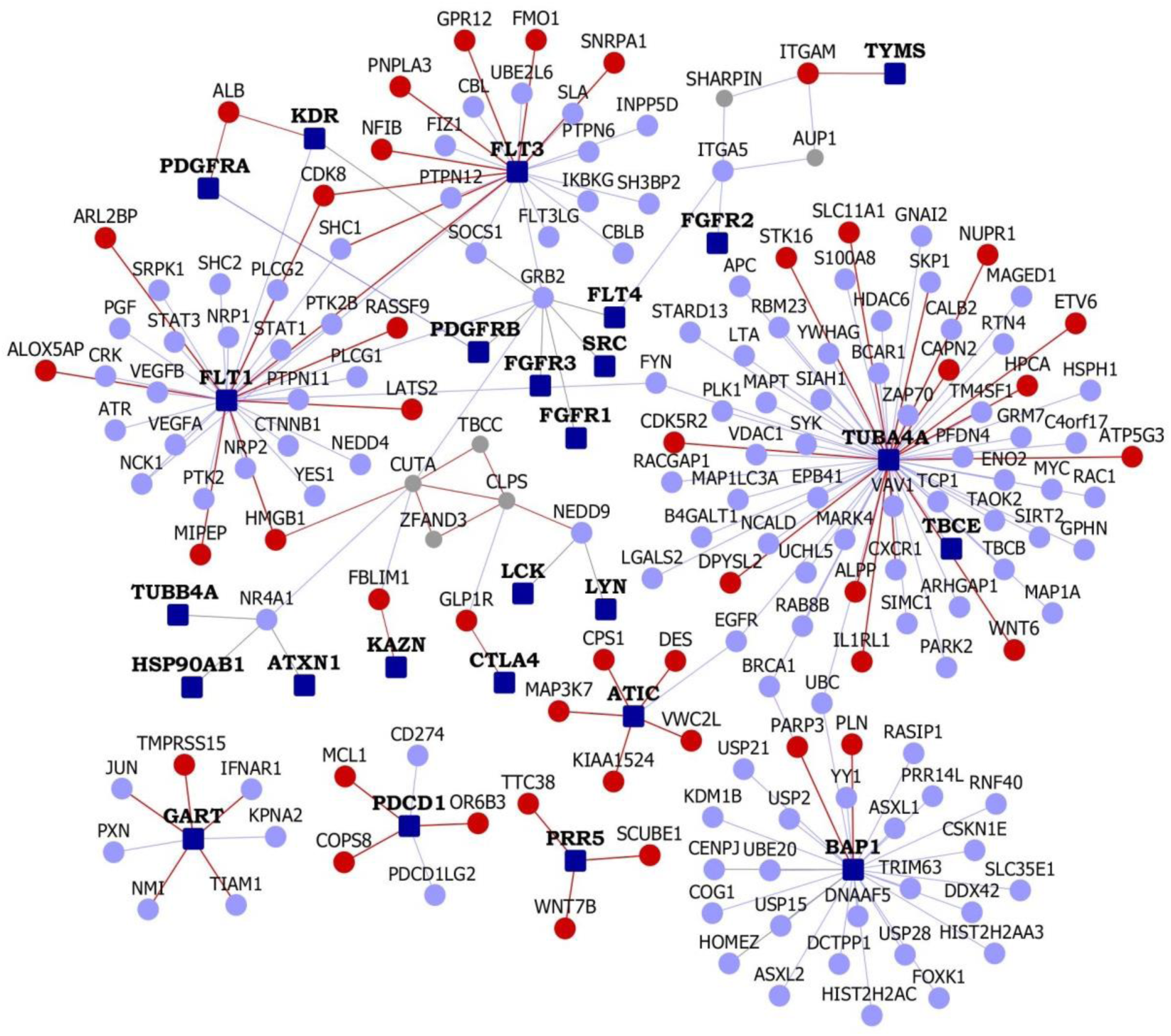
Interactions of selected MPM genes, which are discussed in detail: FLT1, FLT3, TUBA4A, ATIC, CUTA, GART, PDCD1, PRR5 and BAP1. The network around the 5 PPIs (ALB-KDR, ALB-PDGFRA, BAP1-PARP3 and CUTA-CLPS) which have been validated may also be seen.

The work flow illustrated here is applied to other proteins, PPIs and pathways in the MPM interactome to draw the following additional conclusions.

### Novel interactions of FLT genes perturbed on crocidolite exposure may influence angiogenesis and metastasis

The novel interactions of FLT1 and FLT3 may provide insights into various aspects of angiogenesis that affect the pathophysiology of MPM in the context of differentially expression upon asbestos or asbestos-like particle exposure. A study has demonstrated the formation of vascular network or tubules in human endothelial cells on exposure to crocidolite fibers (96). Inhibition of EGFR signaling that regulates angiogenic growth factors resulted in a significant reduction in release of IL-8, VEGFA, VEFGR1 and VEGFR2 (96). Novel interactions of FLT1 with HMGB1 and LATS2, and FLT3 with FMO1 and NFIB may perturb VEFG signaling pathway on crocidolite exposure (Figure 8). LATS2 is a tumor suppressor gene that is inactivated in one-third of mesothelioma cell lines or subjected to copy number deletion on exposure to asbestos fibers (97, 98). MicroRNA miR-93 promotes angiogenesis and metastasis of tumors by suppressing the expression of LATS2, thereby increasing cell survival, tube formation and invasion (99). The effect of LATS2 on angiogenesis and metastasis was previously unexplained.

The novel interaction of LATS2 with FLT1 uncovered in our study opens up the possibility of LATS2 being involved in VEFG signaling. The expression of miR-93 in mouse metastatic model promoted the metastasis of tumor cells to lung tissue (99). miR-93 was also upregulated in non-small cell lung cancer (100). MicroRNAs serve as potential diagnostic and prognostic markers of mesothelioma and efforts have been underway to correlate their expression with asbestos exposure (90, 101). Hyperoxia in tissue has been shown to enhance formation of new blood vessels (102). In NFIB hemizygous mice, the expression of HIF1α, which targets pro-angiogenic factors, was significantly increased under hyperoxia conditions (103).

### Novel interactions of PROTOR may contribute to aberrant activation of PI3K/AKT signaling

The novel interactions of PRR5 (or ‘PROTOR’) with WNT7B, SCUBE1and TTC38, may shed light on the mechanism by which the PI3K/AKT signaling becomes aberrantly active in MPM, which has not been understood before (Figure 8) (56). PRR5 has been implicated in malignant mesothelioma (104). It binds directly to mTOR and RICTOR within the mTORC2 complex which phosphorylates and activates AKT in a TSC1/TSC2 complex-dependent manner, promoting cell growth and proliferation (105, 106). A role for PRR5 in the regulation of RICTOR-mediated recruitment of mTOR substrates or other signaling molecules has been conjectured (107). Silencing of PRR5 inhibits phosphorylation of AKT and S6K1, both of which are downstream targets of mTORC2 (107). Silencing of RICTOR also inhibits AKT activity (107). We hypothesize that the activation of AKT is brought about by WNT7B binding to PRR5, aided by SCUBE1 and TTC38.

It is known that Wnt ligands activate other signaling molecules including mTORC1 and mTORC2 (108, 109). WNT7B, which is implicated in the proliferation of the lung mesenchyme, is also the only Wnt protein that is expressed in the airway epithelium (109, 110). The novel interactor SCUBE1 belongs to a family of secreted proteins involved in inflammatory pathways and organ development, and is implicated in the pathogenesis of lung adenocarcinoma and non-small cell lung cancer (111, 112). SCUBE1 forms a hetero-oligomer with SCUBE2 (112), which promotes AKT activity (113). Proteins that possess tetratricopeptide repeat domains provide binding surfaces for PPIs and also promote the formation of multi-protein complexes (114). So in our case, the novel interaction of PRR5 with TTC38 may be serving to facilitate the formation of such a multi-protein complex that also constitutes WNT7B and SCUBE1. TTC38 was differentially expressed in a spheroid model of mesothelioma (115, 116). TTC38 interacts with TK1 which is involved in non-canonical Wnt pathway (117).

### Novel Interactions of ATIC may underlie malignant transformation of pleural cells aided by biosynthetic-cell survival pathways

Cancerous cells have different metabolic needs compared to normal cells. In order to support over-proliferation of cells, nutrients are consumed excessively and channeled into biosynthetic pathways (118). In such a scenario, many important oncogenic pathways converge to modulate metabolism of tumor cells (119). This convergence may even serve specific purposes over the course of cancer progression such as malignant transformation of tumor cells (Figure 8) (119). ATIC is a bi-functional enzyme that catalyzes the last two steps in biosynthesis of purines in addition to being involved in metabolism of folates (120). CPS1 is a mitochondrial enzyme that catalyzes the first committed step of the urea cycle and is critical in preventing accumulation of toxic ammonia. When CPS1 is knocked down in LKB1-inactivated lung adenocarcinoma cells, there was a reduction in the level of metabolites associated with purine synthesis, and in cell growth (121). LKB1is a master kinase found to be mutated in 20% of non-small cell lung cancers and in mesothelioma cell lines. It regulates the activity of AMPK-related kinases which are similar to AMPK. AMPK acts as a sensor of energy status in cells upon phosphorylation (*122-124*). AMPK blocks growth of cancer cells, and inhibits anabolic processes and instead promotes catabolic processes in order to conserve cellular energy (125). LKB1-mutant cells showed a decrease in AMPK phosphorylation even upon treatment with a known AMPK activator AICAR which is an intermediate of the purine biosynthesis pathway acted upon by ATIC, the rate-limiting enzyme of this pathway (126). Inducing overexpression of CPS1 in LKB1-inactivated lung adenocarcinoma cell lines or CPS1-negative lung adenocarcinoma cell lines did not induce cell growth (121). This suggested that the role played by CPS1in cell growth might be complemented by other genes in CPS1-negative cell lines. CPS1 overexpression in LKB1-inactivated cells failing to induce cell growth may indicate that purine biosynthesis (even though shown to be affected by CPS1 knockdown) was regulated differently in cancer cells (121). The fact that the treatment of LKB1-mutant cells with AICAR failed to activate AMPK, which is necessary for inactivation of the purine biosynthesis pathway and thereby control of cell growth, might also point at a different pathway or set of interactions regulating purine biosynthesis and growth of these cancerous cells (126). Novel interactions of ATIC may throw some light in this direction, viz., a convergence of a metabolic pathway, purine biosynthesis in this case, with oncogenic pathways. ATIC has been predicted to interact with CIP2A. Close correlation has been found between expression levels of CIP2A and c-MYC, an oncogene that promotes cell growth (127). CIP2A inhibits PP2A which can then no longer de-phosphorylate and inactivate members of the MAP kinase family (128). CIP2A is also required for anchorage-independent growth and malignant transformation of human cells (129). A gene-gene interaction (GGI) in which MYC overexpression causes lethality in cells in which CPS1 has been mutated was previously reported (130). This GGI between MYC and CPS1, and the novel PPI between ATIC and CPS1 may be perturbed in MPM. Another novel interactor of ATIC is MAP3K7. Members of the MAP kinase pathway are well-characterized as downstream targets of IRS proteins contributing to malignant phenotype in breast cancer (131). MPM cell lines showed increased MAPK activity in response to IGF1 treatment (132). Another novel interactor of ATIC is DES, a biomarker used to distinguish between malignant mesothelioma and reactive mesothelioma cells, raising the suspicion that these novel interactions may serve malignant transformation of pleural cells (133). A combination of positive epithelial membrane antigen and negative DES as identified from immunohistochemistry is an indicator of malignant mesothelioma (133). VWC2L, another novel interactor of ATIC interacts with CHEK1, a regulator of the cell cycle also involved in breast cancer metastasis and TRAF2, a protein that suppresses death receptor 5 enhancing invasion and metastasis of cancer cells (unpublished AP-MS results obtained from BioGRID) (*134-136*). Since Pemetrexed, a drug used in treatment of mesothelioma that targets ATIC along with other key enzymes involved in synthesis of nucleic acids, interferes with folate metabolism, the involvement of the latter in this convergence of pathways cannot be ruled out (137).

### Novel interactions of FLT genes may underlie induction of angiogenic responses and de-regulation of endocytic receptor trafficking in mesothelioma

Cell survival and growth are intricately linked to inflammatory pathways in pleural mesothelioma (138). Asbestos induces necrosis of cells causing them to release HMGB1, a protein that recognizes danger associated molecular patterns (DAMPs), i.e., signals associated with cellular stress (139). HMGB1 then activates the inflammasome NALP3 that induces pro-inflammatory responses leading to secretion of IL-1β and TNFα and activation of NFκβ, all of which contribute to increase in cell survival and growth (139). Thus, the role of HMGB1 as a pro-inflammatory cytokine has been critical to explaining the pathogenesis of mesothelioma.

The emerging role of HMGB1 as an inducer of angiogenic factors is more interesting in terms of drug development (140–142). Overexpression of HMGB1 increases the angiogenic potential of endothelial cells through stimulation of VEGFR (141). It induced the expression of proangiogenic factors such as VEGFA, and VEGF receptors/co-receptors FLT1, KDR and NRP1, and stimulated abnormalities in angiogenesis at microvascular level, apart from increasing vessel density and dilation (33, 143, 144). Studies reveal a complex interplay between HMGB1 and VEGF signaling in tumor cells and tumor associated macrophages. While it was shown that HMGB1 indirectly influences VEGF receptors, the novel interaction of HMGB1 with FLT1, which has been experimentally validated here, highlights the possibility of a direct connection between this cytokine and the VEGF receptors (Figure 8). HMGB1 expression parallels FGF activation of endothelial cells, and from its known PPIs in the interactome, we could find that the receptors FGFR1, FGFR2 and FGFR3 interacted with VEGF receptors through intermediaries, viz. NCK1 and GRB2 (143). In particular, GRB2 is an adaptor protein, which acts as a critical link between growth factor receptors and the Ras signaling pathway, and is also involved in downstream signaling of FGFR2 (145). NCK1 is another adaptor protein that associates with growth factor receptors such as KDR or their cellular substrates (146). HMGB1 is reported to interact with TLR4 receptor to mediate pro-inflammatory responses (147). In the interactome, TLR4 was found to interact with HSP90B1 that participates in VEGF signaling by stabilizing and folding other proteins (148). It appears that the receptor activity of TLR4 and its indirect interaction with FLT1 may be mediated by HSP90B1 and some kinases including SRC. It is known that HMGB1 increases permeability of endothelial cells through SRC family tyrosine kinases (149).

The novel interaction of both the receptors FLT1 and FLT3 with CDK8 may throw some light into the manner in which expression of proangiogenic factors in the VEGF signaling pathway may be regulated (Figure 8). HIF1A employs CDK8 for its downstream activities and overexpression of HIF1A has been correlated with increased expression of its target genes, viz. proangiogenic factors of the VEGF signaling pathway (150). In the interactome, HIF1A was found to interact with FLT1 through HSP90AA1 that participates in VEGF signaling and mediates proper folding of target proteins aided by other co-chaperones (148). A novel interaction between the receptors FLT1 and FLT3 was also found (Figure 8). Such receptor-receptor interactions are not without precedence – the orphan receptor kinase TIE1, involved in vascular development, was identified in endothelial cells to bind to TEK (151). It has been speculated that receptors that are physiologically related, such as VEGF receptors, may display weak interactions among their transmembrane domains (152).

VEGF signaling pathway is tightly regulated by processes such as receptor endocytosis, internalization and intracellular trafficking (153). Endosomes play a critical role in these processes by allowing signaling from their compartments, serving as scaffolds to facilitate the assembly and degradation of signaling complexes, and trafficking receptors to many subcellular locations (153). For example, VEGFR2 is rapidly internalized to endosomes and degraded on binding of VEGF (153). Similarly, the assembly of VEGFR2 is contingent upon endocytic trafficking (153). We think that the novel interactions of RASSF9 and ARL2BP with FLT1 may be relevant in this respect (Figure 8). RASSF9 is an endosomal protein that helps in the trafficking of PAM through the secretory or endosomal pathways (154). ARL2BP is a regulator of vesicle formation during intracellular trafficking and was found to be overexpressed in conditions of TLR4 dependent inflammation and tumorigenesis (155). Moreover, crocidolite fibers that induce changes in the expression pattern of HMGB1, a ligand of the TLR4 receptor, are actively transported in endosomes to secondary lysosomes located near the nucleus in a size-dependent manner in epithelial cells of lung (156).

### Novel interactions of IFNAR1 and GART may point at the influence of immune system over metastatic pathways converging to serve tumor metabolism in mesothelioma

During the different stages of cancer progression, tumor cells adopt a number of strategies to evade surveillance by immune cells (157). Pleural effusions containing malignant mesothelial cells are replete with a wide range of immune cells that have infiltrated these tumors; yet, the immunological status of mesothelioma patients is tolerant towards these cancerous cells (158). The complex interplay of tumor-immune interactions enables tumors to evade immune cells. Some of the novel interactions in the MPM interactome may throw light on immunological pathways that are deregulated in cancer to aid processes such as tumor invasion and metastasis.

In mice lacking IFNAR1, loss of type-1 interferon signaling is sufficient to promote metastasis of breast cancer to lung, independently of the growth of primary tumors (159). Reduced expression of CD69 and IFNγ, two primary biomarkers of Natural Killer (NK) cells, also showed that the homeostasis of the NK population in the immune system which is dependent on signaling through IFNAR1 receptors was impaired in the process (159). The cytokine IL-2 is used as an immunotherapeutic agent to induce anti-metastatic and cytotoxic effects in cancer cells and has been approved for the treatment of metastatic renal cell carcinoma and melanoma. In mice lacking IFNAR1 receptors, immunotherapy using IL-2 was unsuccessful. This showed that the cytotoxic effect mediated by this cytokine was abolished in the absence of IFNAR1 signaling (160). The interleukin-2 receptor, IL2Rβ, was found to be significantly downregulated in a small-cell lung cancer cell line, compared with normal lung tissue (SF Table 3). The exact mechanism underlying this influence of the immune system over metastatic process was unknown. Novel interaction of IFNAR1 with GART and novel interactions of GART- a protein involved in *de novo* purine biosynthesis- with a host of proteins that regulate tumor invasion viz. TIAM1, NMI, PXN, KPNA2, TMPRSS15 and JUN may throw some light in this direction (Figure 8). The said novel interactions may depict a convergence of many oncogenic pathways to modulate metabolism of tumor cells and serve specific purposes, viz. malignant transformation of tumor cells, as has been mentioned previously for the novel interactions of ATIC. However, this convergence may also be immunologically regulated by NK-cell receptors, viz. KIR2DL3 which interacts with LCK, a known interactor of IFNAR1, and novel interactors of PDCD1 such as MCL1 and COPS8 (Figure 8) potentially serving as downstream targets (probably influenced by DHFR and its novel interactor ASGR1) with HLA-DQA1 as the ligand on the metastatic cell.

In non-small cell lung cancer, a higher proportion of NK cells express KIR2DL3, which can be induced by IL-2 and reduces with advancing disease (161, 162). LCK belongs to the family of SRC kinases that regulate cell proliferation, differentiation, survival and cytoskeletal alterations (163). HLA-DQA1 is significantly downregulated in small cell lung cancer cell lines compared to their expression in control tissue samples of lung or lymph node (SF Table 3). Hence, these novel interactions may point to NK-tumor cell interactions in metastatic locations of the lung or lymph nodes critical to the progression of MPM.

High expression of GART is observed in cell lines of non-small cell lung cancer; GART and other genes involved in purine biosynthesis such as PPAT, PAICS, PKM2 and ATIC, have been positively correlated with increased proliferation of lung cancer cells (164). One of the novel interactors of GART is a type II transmembrane serine protease that takes part in epithelial-mesenchymal transitions called TMPRSS15, the activity of which is dysregulated in cancer in order to degrade and remodel intercellular junctions and the extracellular matrix (165). Silencing of TMPRSS15 leads to tumor migration and matrix degradation in lung cancer cell lines (165). NMI, another novel interactor of GART, negatively regulates epithelial-mesenchymal transitions by inhibiting p65 acetylation in the NF-Kβ pathway (166). Knockdown of KPNA2, a novel interactor of GART, suppresses the proliferative and migratory abilities of lung cancer cells and downregulation of its regulatory miRNA miR-26b enhances cell migration and invasion in vitro, and metastasis in vivo (167). One of the downstream targets of KPNA2, which is overexpressed in non-small cell lung cancer, is another novel interactor of GART called JUN, a component of the transcription factor AP-1 promoting progression of cell cycle and involved in breakdown of the ECM (168). TIAM1, a novel interactor, is a guanosine exchange factor that takes part in the activation of c-Jun-terminal kinase, p38 MAPK and ERKs, and regulation of genes involved in cellular migration, invasion and metastases (169). PXN, a novel interactor of GART, is a cytoskeletal protein involved in membrane attachment of actin at sites of cell adhesion to extra-cellular matrix (169). Mutations in PXN are associated with lung adenocarcinoma and its overexpression induced by suppression of miR-218 is correlated with increased cell proliferation and invasion (169). Overexpression of PDCD1, playing a key role in immune evasion and the formation of tumor microenvironment has been correlated with poor prognosis and high invasiveness in non-small cell lung cancer (170). MCL1, a novel interactor of PDCD1, is highly expressed in NK cells and upregulated by IL-15 through IL-2Rβ/γ (which is also a receptor of IL-2) (171). Post mortem analyses of mice having MCL1 deleted and injected with murine melanoma cells revealed overwhelming spread of melanoma cells to lung (171). miR-146a that targets COPS8, another novel interactor of PDCD1, suppresses the expression of tumor promoting cytokines and growth factors in gastric adenocarcinoma (172). DHFR, which is involved in folate metabolism and cell growth, has also been predicted to interact with COPS8 (173). Folate metabolism is critical to cancer development as is indicated by the efficacy of anti-folates such as Pemetrexed in treatment of cancer (174). Pemetrexed, a drug used for treatment of MPM in combination with Cisplatin, targets DHFR (62).

Both mesothelioma and lung cancers share common pathways that are initiated on exposure to asbestos fibers in mesothelial cells and lung epithelial cells respectively (175). So, RNA-Sequencing data obtained from pathology atlas for normal lung or lymph tissue were compared with data from the small cell lung cancer cell line, SCLC-21H. It was observed that all the genes that perform anti-metastatic functions in normal tissue were down regulated in SCLC-21H, viz. IFNAR1, LCK, HLA-DQA1 and NMI (SF Table 3). On the other hand, genes that perform pro-metastatic functions, viz. KPNA2, GART, PDCD1 and COPS8 were found to be upregulated (SF Table 3). On comparing RNA-sequencing data obtained from GSE9586 it was found that the natural killer receptor KIR2DL3 was significantly down regulated under conditions favorable to metastasis, which in this case is a metastatic breast cancer cell line in which the microRNA miR-335 is not expressed (Log2FC = −2.659, P-value = 0.018). miR-335 inhibits metastatic cell invasion by regulating a set of genes in tumors that are associated with risk of metastasis to distant sites (176).

### Limitations of results and interpretations

It is beyond the scope of our expertise to validate the large number of computationally predicted PPIs in a tissue or cell line of interest. However, we demonstrated the validity of computational predictions on a small number of PPIs on purified proteins with appropriate controls. The computational model has also been validated through additional experiments previously and the novel PPIs predicted in other contexts have translated into results of biomedical significance (*19-21*). This will catalyze further investigations into these particular PPIs and may lead to biologically or clinically translatable results.

Secondly, biologists often formulate hypotheses around individual ‘components’ of a system, for example, a single protein or a pathway based on their knowledge and by studying existing databases and literature, which are then validated through resource-intensive and time-consuming experiments. In this paper, we presented a large set of PPIs relevant to MPM. However, by making these results available on a searchable web database, we enable biologists to choose the PPI of interest from the MPM interactome by querying for it (http://severus.dbmi.pitt.edu/wiki-MPM). Our website provides advanced search capabilities which allows a user to ask questions such as “show me the PPIs in which one protein is involved in mesothelioma and the other is involved in inflammation” and then see the results with the functional details of the two proteins side-by-side. This will help biologists to generate testable hypotheses around individual PPIs and advance the biology surrounding each PPI by performing suitable experiments. A further advantage is that these resources will also be indexed in popular internet search engines like Google and Bing, and the novel and known PPIs relevant to mesothelioma would also be found through them.

### Translation of results for clinical applications

We presented 5 drugs, namely *cabazitaxel, primaquine, pyrimethamine, trimethoprim and gliclazide*, which may potentially be repurposed for treating MPM. Preclinical studies may be conducted *in vitro* to validate these computational results.

## Conclusions

Diagnosis of mesothelioma, which has a long latency period of more than 30 years, is difficult due to a lack of distinctive symptoms in patients. It recurs after surgical procedures such as extrapleural pneumonectomy, radiation therapy and post-surgical chemotherapy (177). Moreover, the median survival time for mesothelioma is 12.1 months after first-line therapy with pemetrexed/cisplatin. An increase in its incidence has been predicted in western countries and economically emerging nations based on studies on asbestos utilization (177). In this scenario, it is essential that we accelerate biomedical discovery in this field, to catalyze the emergence of clinically translatable results. In this paper, we show that interactome-level analysis may be a right step in this direction. The MPM interactome with MPM-associated proteins and their interacting partners will help biologists, bioinformaticians and clinicians to piece together an integrated view on how genes associated with MPM through various high throughput studies are functionally linked. Such biological insights will lead to clinically translatable results such as testable hypotheses centered on individual protein-protein interactions based on pathway analysis, and drugs repurposable for mesothelioma.

## Materials and Methods

### Data collection

Genes associated with malignant pleural mesothelioma were obtained from IPA (Ingenuity Pathway Analysis) (178). A search on IPA using the keyword “malignant pleural mesothelioma” retrieved genes that are causally relevant to the disease. The genes are retrieved from the Ingenuity Knowledge Base, a structured collection of nearly 5 million findings with experimental basis which have been manually curated from biomedical literature or incorporated from other databases (178).

### Prediction model-HiPPIP

PPIs were predicted by computing features of protein pairs and developing a random forest model to classify the pairwise features as *interacting* or *non-interacting*. Protein annotations that were used in this work are: cellular localization, molecular function and biological process membership, location of the gene on the genome, gene expression in hundreds of microarray experiments, protein domains and tissue membership of proteins. Computation of features of protein-pairs is described earlier in Thahir et al (28). A random forest with 30 trees was trained using the feature offering maximum information gain out of 4 random features to split each node; minimum number of samples in each leaf node was set to be 10. The random forest outputs a continuous valued score in the range of [0,1]. The threshold to assign a final label was varied over the range of the score for positive class (i.e., 0 to 1) to find the precision and recall combinations that are observed. This prediction model is referred to as High-confidence Protein-Protein Interaction Prediction (HiPPIP) model.

### Evaluation of PPI prediction model

Evaluations on a held-out test data showed a precision of 97.5% and a recall of 5% at a threshold of 0.75 on the output score. Next, we created ranked lists for each of the hub genes (i.e., genes that had more than 50 known PPIs), where we considered all pairs that received a score greater than 0.5 to be novel interactions. The predicted interactions of each of the hub genes are arranged in descending order of the prediction score, and precision versus recall is computed by varying the threshold of predicted score from 1 to 0. Next, by scanning these ranked lists from top to bottom, the number of true positives versus false positives was computed.

### Novel PPIs in the MPM interactome

Each MPM gene, say Z, is paired with each of the other human genes (G_1_, G_2_ … G_N_), and each pair is evaluated with the HiPPIP model. The predicted interactions of each of the MPM genes (namely, the pairs whose score is greater than the threshold 0.5) were extracted. These PPIs, combined with the previously known PPIs of MPM genes collectively form the *MPM interactome.* Interactome figures were created using Cytoscape (179).

Note that 0.5 is the threshold chosen not because it is the midpoint between the two classes, but because the evaluations with hub proteins showed that the pairs that received a score greater than 0.5 are highly confident to be interacting pairs. This aspect was further validated by experimentally validating a few novel PPIs above this score.

### In vitro pull-down assays

An in vitro pull-down assay method was employed to determine physical interactions between proteins and to perform initial screening of some of the predicted novel protein-protein interactions. This technique relies on utilizing a tag-fused protein (e.g., His-tag, biotin-tag) immobilized on an affinity column or a resin as the bait protein and a passing-through solution containing the ‘prey’ protein that binds to the ‘bait’ protein. The subsequent elution will pull down both the target (prey) and tagged-protein (bait) for further analysis by immunoblotting to confirm the predicted interactions. The pull-down assays to validate some of the chosen predicted interactions were conducted using the Pull-Down PolyHis Protein:Protein Interaction Kit (Pierce™) according to the manufacturer’s instructions. Briefly, the His-tagged bait proteins (i.e., CLPS, HMGB1, VEGFR2/KDR, PDGFRA) and untagged prey proteins (i.e., CUTA and ALB), purchased from either MyBiosource or Abcam, were diluted to desired concentrations (50 μg/ml) in the tris-buffered saline solution (TBS; 25mM Tris·HCl, 0.15M NaCl; pH 7.2). 200 μl of TBS solution containing each bait protein was incubated with 25 μl of Cobalt Resin (HisPur™; Pierce) at room temperature for 30 min to capture the His-tagged bait protein. The beads were then washed 5x with 200 jliI of wash buffer (TBS containing lOmM Imidazole) and by centrifuging at 1250 x g for 1 minute to remove unbound bait protein. 200 μl of TBS buffer containing the desired untagged prey protein was then added to the above bait containing beads. This mixture was incubated at RT for ~1 hour. Following this, the beads were washed thrice in the same manner described above and finally the bound complex in each case was eluted using 300 μl of elution buffer (wash buffer containing 290mM Imidazole). Thus eluted samples in each case were concentrated to ~50ul and were further analyzed using Wes™ Simple Western, a capillary western blot technology which separates and analyzes proteins by size from 2-440 kDa. A total protein simple western size-based assay was performed according to the ProteinSimple user manual. In brief, 4ul of concentrated samples from the elution’s of pull-down assay were mixed with a master mix (ProteinSimple, Santa Clara, CA) to a final concentration of 1×sample buffer, 1 × fluorescent molecular weight markers and 40mM DTT. Thus prepared protein samples were then heated at 95 °C for 5 minutes before loading on to the plate. These samples along with biotinylated ladder, diluent, biotin labeling reagent, streptavidin-HRP, wash buffer and luminol-peroxide mix were also dispensed in to corresponding wells as indicated in the plate diagram. Thus prepared plate was centrifuged at RT for 5mins @ 2500 rpm. The run was carried out at room temperature using instrument’s default settings. The chemiluminescence generated was captured by a CCD camera. The digital image was analyzed with Compass software (ProteinSimple).

The pseudo-gel and electro-pherogram profiles, showing bands and peaks corresponding to both ‘bait’ and ‘prey’ proteins in the pull-down samples, highlight and suggest the potential for the interaction of ALB with KDR/VEGFR2 and PDGFRA (Figure 9). It has to be noted that some of the selected proteins under reducing conditions may exhibit higher apparent mass than absolute molecular weights, due to post-translational modifications (e.g., KDR and PDGFRA) or in the presence of metals (e.g., CUTA) (180). In the case of CUTA interactions, a band or peak around 74kDa was observed in the washes after CUTA protein binding as ‘prey’ (Figure 2, lane 6/9). It has been previously reported that CUTA can form multimers under certain experimental conditions including the presence of metals/metal ions. The monomeric form of CUTA migrates at 17-18kDa, CLPS at 13kDa and HMGB1 migrates at 28 kDa and 32 kDa. While HMGB1-CUTA pull-down sample resulted in a broad band/peak around 74-80kDa and 36-52kDa, CLPS-CUTA showed a broad band/peak around 13-24KDa along with several minor peaks at 34kDa, 74kDa and 115kDa. This may suggests a strong interaction of CLPS with CUTA, which interferes with the agglomeration state of CUTA. This is further supported by the fact that the intensity of the bands and peak height corresponding to CUTA in the wash after binding to CLPS as a ‘bait’ is less or almost non-existent compared to wash after adding to HMGB1. CUTA-CLPS was also validated through LC-MS (SF Table 1).

**Figure 9.**
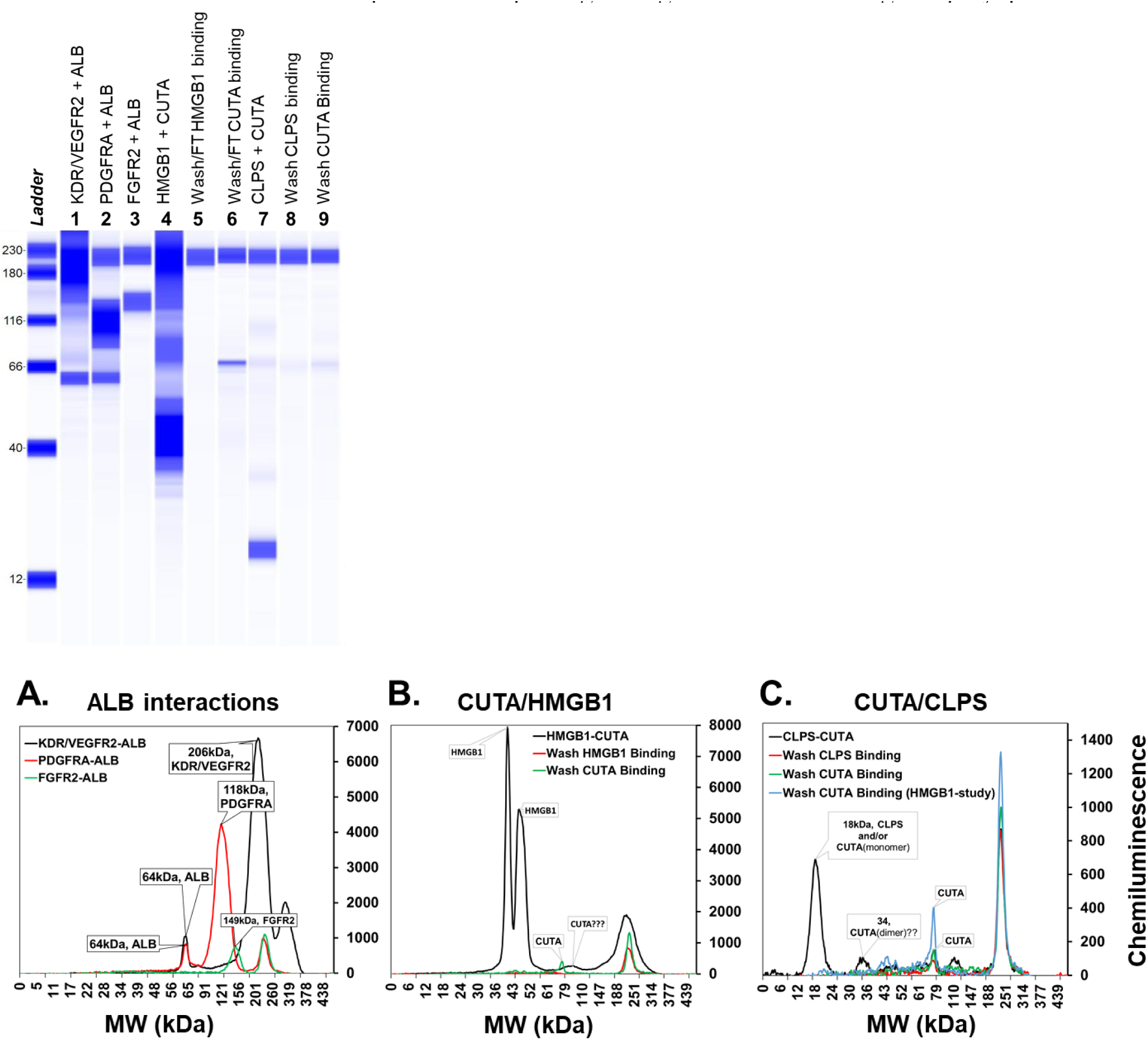
Validation of predicted ALB interactions and CUTA interactions using Wes^™^ Simple Western total protein detection assay. Pseudo-gel or virtual-blot like image of the validated interactions of ALB (lanes 1-2) and CUTA (lanes 4, 7) along with negative control (lane 3). In addition to the final pull-down samples, wash and/or flow through after binding ‘bait’ and ‘prey’ proteins for the CUTA interactions are also shown (lanes 5,6,8 & 9). The electro-pherogram image of Simple Western results using Total protein size-based assay. (A) ALB interactions with true positives KDR/VEGFR2, PDGFRA and false positive FGFR2. (B) CUTA interactions with HMGB1. (C) CUTA interactions with CLPS. An overlay of the electro-pherogram of the wash from HMGB1 after CUTA binding is also shown in (C) for comparison.

### Protein identification methods

Peptide sequencing experiments were performed using an EASY-nLC 1000 coupled to a Q Exactive Orbitrap Mass Spectrometer (Thermo Scientific, San Jose, CA) operating in positive ion mode. An EasySpray C18 column (2 μm particle size, 75 μm diameter by 15cm length) was loaded with 500 ng of protein digest in 22 μL of solvent A (water, 0.1% formic acid) at a pressure of 800 bar. Separations were performed using a linear gradient ramping from 5% solvent B (75% acetonitrile, 25% water, 0.1% formic acid) to 30% solvent B over 120 minutes, flowing at 300 nL/min.

The mass spectrometer was operated in data-dependent acquisition mode. Precursor scans were acquired at 70,000 resolution over 300-1750 m/z mass range (3e6 AGC target, 20 ms maximum injection time). Tandem MS spectra were acquired using HCD of the top 10 most abundant precursor ions at 17,500 resolution (NCE 28, 1e5 AGC target, 60 ms maximum injection time, 2.0 m/z isolation window). Charge states 1, 6-8 and higher were excluded for fragmentation and dynamic exclusion was set to 20.0 s.

Mass spectra were searched for peptide identifications using Proteome Discoverer 2.1 (Thermo Scientific) using the Sequest HT and MSAmanda algorithms, peptide spectral matches were validated using Percolator (target FDR 1%). Initial searches were performed against the complete UniProt database (downloaded 19 March, 2018). Peptide matches were restricted to 10 ppm MS1 tolerance, 20 mmu MS2 tolerance, and 2 missed tryptic cleavages. Fixed modifications were limited to cysteine carbamidomethylation, and dynamic modifications were methionine oxidation and protein N-terminal acetylation. Peptide and protein grouping and results validation was performed using Scaffold 4.8.4 (Proteome Software, Portland, OR) along with the X! Tandem algorithm against the previously described database. Proteins were filtered using a 99% FDR threshold.

### Differential gene expression in the small cell lung cancer cell line SCLC-21H

Gene expression data of the small-cell lung cancer cell line, SCLC-21H, and normal lung tissue were obtained from Human Cell Atlas(181) and Human Tissue Atlas(182) respectively. For each gene, the expression values were duplicated in SCLC-21H, and replicated nine times in the normal lung tissue. The deviation of the gene expression values of the different replicates from the mean value was calculated using 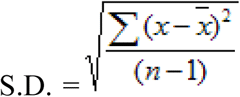 where x̅ is the mean average of the replicated gene expression values within a group (i.e. within the test or control group) and n is the sample size.

P-value of significance of the observed difference between the means of the test and control groups was calculated using the t-test, with the value t calculated as 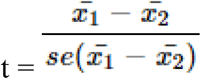, where x̅ is the mean average and se is the standard error of the difference between the means of the test and control groups.

The fold change in expression for each gene was calculated as the ratio of the average TPM (transcripts per million) value of the gene in SCLC-21H (test) and the corresponding value in normal lung tissue (control). Genes with fold change >2 or ½ were considered as significantly overexpressed and underexpressed respectively at P-value<0.05.

### Analysis of DNA methylation in MPM tumors

The dataset GSE16559 (42) deposited in GEO was used to analyze the methylation profile of pleural mesotheliomas. In this study, genes found to be differentially methylated in mesothelioma were identified from a set of 773 cancer-related genes associated with 1413 autosomal CpG loci. Methylation values (or M-values) were computed as M = log2 [β (1-β)] for both control (non-tumor pleural tissue) and test (pleural mesothelioma) cases, where β is the ratio of methylated probe intensity and overall intensity. Difference between M-values of test and control cases was then computed and genes with M-value>1 and M-value<1 were considered to be hypermethylated and hypomethylated respectively at P-value<0.05.

### Analysis of differential gene expression in pleural mesothelioma tumors and normal tissue in lungs

The overlap of the MPM interactome with genes differentially expressed in pleural mesothelioma tumors compared with normal pleural tissue adjacent to mesothelioma was computed using the dataset GSE12345(48). A total of 22876 were assayed in this dataset, out of which 3162 were differentially expressed in pleural mesothelioma tumors. 142 genes in the MPM interactome were not assayed in this dataset and for computing the overlap, this was added to the set of genes assayed in the expression study, giving a total of 22918 genes. The overlap of the MPM interactome with genes that distinguish MPM tumors from other thoracic cancers was computed using the dataset GSE42977 (32) containing the expression of MPM tumors vs. other thoracic cancers such as thymoma and thyroid cancer. A total of 24996 genes were assayed in this dataset, out of which 6176 were differentially expressed in MPM tumors. 222 genes in the MPM interactome were not assayed in this dataset and for computing the overlap, this was added to the set of genes assayed in the expression study, giving a total of 25218 genes. Genes with fold-change >2 or <½ was considered as overexpressed and under expressed respectively at P-value<0.05. Genes which have high/medium expression in normal lung tissue (median TPM≥9), were identified using RNA-sequencing data available in GTEx (49).

### Correlating expression of MPM genes with lung cancer prognosis

Data for correlation of gene expression and fraction of patient population surviving after treatment for lung cancer was taken from Pathology Atlas (50). A total of 19622 genes were assayed in this dataset, out of which the expression of 354 genes were correlated with unfavorable prognosis. Log-rank P-values which indicate the significance of this correlation were examined. Genes with log-rank P-value<0.001 were considered to be prognostic. Unfavorable prognosis indicates positive correlation of high gene expression with reduced patient survival.

### Interactome of MPM genes differentially expressed on particle exposure

Genes differentially expressed in the lungs of mice exposed to crocidolite and erionite fibers were obtained from the dataset GSE100900(54). A total of 24434 genes were assayed in this dataset, out of which 1710 were differentially expressed on crocidolite exposure. Those genes that were differentially expressed on crocidolite and erionite exposure and their interactions with MPM genes were selected from the MPM interactome. This network was further extended to show how these genes interact with other known genes in the MPM interactome. Interactome figures were created using Cytoscape (179).

### Identification of repurposable drugs in the MPM drug-protein interactome

To identify repurposable drugs already tested in non-small cell lung cancer (NSCLC), drugs tested in (completed) clinical trials of NSCLC were obtained from NIH Clinical Trials (https://clinicaltrials.gov/). Then, the list of drugs targeting MPM genes that had negative correlation with lung cancer expression studies was compared with drugs tested in NSCLC to identify overlaps. Drugs potentially repurposable for MPM were chosen from the list of overlapping drugs based on literature review.

To identify repurposable drugs targeting MPM genes or novel interactors, drugs targeting MPM genes or novel interactors that had negative correlation with lung cancer expression studies were identified. Next, the genes targeted by these drugs were chosen and their differential expression in MPM tumour/cell lines (GSE51024 (60) and GSE2549(35)) was checked. Genes with high fold changes and low P-values were chosen and literature review was conducted to check whether their corresponding drugs were potentially repurposable.

To identify repurposable drugs targeting known interactors, those known interactors which had high fold changes in MPM tumours/cell lines (GSE51024 (60) and GSE2549 (35)) were chosen. Drugs targeting these known interactors, that had negative correlation with lung cancer expression studies, were identified. Literature review was conducted to check whether these drugs were potentially repurposable.

Negative correlation between lung cancer and drugs were studied using the BaseSpace correlation software (https://www.nextbio.com), which uses a non-parametric rank-based approach to compute the extent of enrichment of a particular set of genes (or ‘bioset’) in another set of genes (59).

## Supplementary Materials

Supplementary Methods: Additional details of methods and additional tables.

Data File 1: List of MPM genes and the source of information for their association to MPM, as provided by Ingenuity Pathway Analysis (IPA) suite.

Data File 2: List of genes from the MPM interactome, with their labels (namely, MPM genes, known interactors and novel interactors).

Data File 3: Master table of all biological evidences (gene expressions, genetic variants, methylations) for each of the MPM interactome genes discussed in the paper.

Data File 3: List of all the pathways associated with at least one of the MPM genes, along with their statistical significance of association (with Bonferroni correction).

Data File 4: List of all the drugs that target any of the genes from the MPM interactome, along with their ATC (anatomical, therapeutic and chemical) codes.

## Acknowledgments

We thank Prof. Michael Becich (PI, National Mesothelioma Virtual Bank) for support and for funding for MKG and for comments on the manuscript. We thank Prof. N. Balakrishnan for support and for funding for KBK. MKG thanks Dr. David Boone (Department of Biomedical Informatics) and Dr. J. Richard Chaillet (Office of Research Health Sciences) of University of Pittsburgh for detailed and valuable feedback on the writing of the manuscript. MKG thanks Sai Supreetha Varanasi for system administration assistance in hosting the website.

## Author contributions

In sequence of work: MKG conceptualized and supervised the study and carried out interactome construction and analysis of pathway and drug associations. KBK carried out studies of overlap of the interactome with various high-throughput data, literature-based evidence gathering, and formulation of biological hypotheses. Experimental validations were carried out by NY and GB. Methods of experimental validation were provided by NY and GB. Manuscript has been written by KBK and edited by MKG. Manuscript has been read and approved by all authors.

## Data and materials availability

On journal/preprint archive website and at http://severus.dbmi.pitt.edu/wiki-MPM.

## References

1. S. E. Mutsaers, The mesothelial cell. The international journal of biochemistry & cell biology 36, 9–16 (2004).

2. Z. J. Wang et al., Malignant pleural mesothelioma: evaluation with CT, MR imaging, and PET. Radiographics 24, 105–119 (2004).

3. B. M. Robinson, Malignant pleural mesothelioma: an epidemiological perspective. Annals of cardiothoracic surgery 1, 491 (2012).

4. C. J. Jennings, P. M. Walsh, S. Deady, B. J. Harvey, W. Thomas, Malignant pleural mesothelioma incidence and survival in the Republic of Ireland 1994-2009. Cancer epidemiology 38, 35–41 (2014).

5. C. Norbet, A. Joseph, S. S. Rossi, S. Bhalla, F. R. Gutierrez, Asbestos-related lung disease: a pictorial review. Current problems in diagnostic radiology 44, 371–382 (2015).

6. R. Bueno et al., Comprehensive genomic analysis of malignant pleural mesothelioma identifies recurrent mutations, gene fusions and splicing alterations. Nature genetics 48, 407–420 (2016).

7. J. R. Testa et al., Germline BAP1 mutations predispose to malignant mesothelioma. Nat Genet 43, 1022–1025 (2011).

8. J. A. Ohar et al., Germline BAP1 Mutational Landscape of Asbestos-Exposed Malignant Mesothelioma Patients with Family History of Cancer. Cancer research 76, 206–215 (2016).

9. A. Bononi et al., BAP1 regulates IP3R3-mediated Ca2+ flux to mitochondria suppressing cell transformation. Nature, (2017).

10. R. G. Van Der Most, A. J. Currie, B. Robinson, R. A. Lake, Decoding dangerous death: how cytotoxic chemotherapy invokes inflammation, immunity or nothing at all. Cell death and differentiation 15, 13 (2008).

11. V. Panou et al., Frequency of Germline Mutations in Cancer Susceptibility Genes in Malignant Mesothelioma. Journal of Clinical Oncology, JCO. 2018.2078. 5204 (2018).

12. S. J. Weiner, S. Neragi-Miandoab, Pathogenesis of malignant pleural mesothelioma and the role of environmental and genetic factors. Journal of cancer research and clinical oncology 135, 15–27 (2009).

13. N. Sahni et al., Widespread macromolecular interaction perturbations in human genetic disorders. Cell 161, 647–660 (2015).

14. D. E. Jensen et al., BAP1: a novel ubiquitin hydrolase which binds to the BRCA1 RING finger and enhances BRCA1-mediated cell growth suppression. Oncogene 16, 1097 (1998).

15. A. Kamiya et al., DISC1-NDEL1/NUDEL protein interaction, an essential component for neurite outgrowth, is modulated by genetic variations of DISC1. Human molecular genetics 15, 3313–3323 (2006).

16. G. Griffiths, D. Fennell, A. Casbard, L. Nixon, J. Lester, Vinorelbine in mesothelioma (VIM): a randomised phase II trial of oral vinorelbine as second-line therapy for patients with malignant pleural mesothelioma (MPM) expressing BRCA1-a study in progress. (2012).

17. T. Rolland et al., A proteome-scale map of the human interactome network. Cell 159, 1212–1226 (2014).

18. M. K. Ganapathiraju et al., Schizophrenia interactome with 504 novel protein–protein interactions. npj Schizophrenia 2, 16012 (2016).

19. Y. Li et al., Global genetic analysis in mice unveils central role for cilia in congenital heart disease. Nature 521, 520 (2015).

20. X. Liu et al., The complex genetics of hypoplastic left heart syndrome. Nature genetics 49, 1152 (2017).

21. J. Zhu et al., Antiviral activity of human OASL protein is mediated by enhancing signaling of the RIG-I RNA sensor. Immunity 40, 936–948 (2014).

22. N. Orii, M. K. Ganapathiraju, Wiki-pi: a web-server of annotated human protein-protein interactions to aid in discovery of protein function. PloS one 7, e49029 (2012).

23. S. Nataf, M. Barritault, L. Pays, A Unique TGFB1-Driven Genomic Program Links Astrocytosis, Low-Grade Inflammation and Partial Demyelination in Spinal Cord Periplaques from Progressive Multiple Sclerosis Patients. International journal of molecular sciences 18, 2097 (2017).

24. H. J. Choo, A. Cutler, F. Rother, M. Bader, G. K. Pavlath, Karyopherin alpha 1 regulates satellite cell proliferation and survival by modulating nuclear import. Stem Cells 34, 2784–2797 (2016).

25. S. Cedres et al., Exploratory analysis of activation of PTEN–PI3K pathway and downstream proteins in malignant pleural mesothelioma (MPM). Lung Cancer 77, 192–198 (2012).

26. T. Keshava Prasad et al., Human protein reference database—2009 update. Nucleic acids research 37, D767–D772 (2008).

27. C. Stark et al., BioGRID: a general repository for interaction datasets. Nucleic acids research 34, D535–D539 (2006).

28. M. Thahir, T. Sharma, M. K. Ganapathiraju, An efficient heuristic method for active feature acquisition and its application to protein-protein interaction prediction. BMC proceedings 6 Suppl 7, S2 (2012).

29. M. Cigognetti et al., BAP1 (BRCA1-associated protein 1) is a highly specific marker for differentiating mesothelioma from reactive mesothelial proliferations. Modern Pathology 28, 1043 (2015).

30. B. Lupo, L. Trusolino, Inhibition of poly (ADP-ribosyl) ation in cancer: old and new paradigms revisited. Biochimica et Biophysica Acta (BBA)-Reviews on Cancer 1846, 201–215 (2014).

31. M. Nasu, UNIVERSITY OF HAWAI’I AT MANOA, (2012).

32. A. De Rienzo et al., Sequential binary gene ratio tests define a novel molecular diagnostic strategy for malignant pleural mesothelioma. Clinical Cancer Research 19, 2493–2502 (2013).

33. Z.-H. Yao et al., Serum albumin as a significant prognostic factor in patients with malignant pleural mesothelioma. Tumor Biology 35, 6839–6845 (2014).

34. M. L. Iacono et al., Targeted next-generation sequencing of cancer genes in advanced stage malignant pleural mesothelioma: a retrospective study. Journal of thoracic oncology 10, 492–499 (2015).

35. G. J. Gordon et al., Identification of novel candidate oncogenes and tumor suppressors in malignant pleural mesothelioma using large-scale transcriptional profiling. The American journal of pathology 166, 1827–1840 (2005).

36. Y. Liu, X. Zou, G. Sun, Y. Bao, Codonopsis lanceolata polysaccharide CLPS inhibits melanoma metastasis via regulating integrin signaling. International journal of biological macromolecules 103, 435-440 (2017).

37. T. Okamoto et al., CD9 negatively regulates CD26 expression and inhibits CD26-mediated enhancement of invasive potential of malignant mesothelioma cells. PloS one 9, e86671 (2014).

38. S. Pereira, C. Lowell, The Lyn tyrosine kinase negatively regulates neutrophil integrin signaling. The Journal of Immunology 171, 1319–1327 (2003).

39. V. V. Orlova et al., A novel pathway of HMGB1-mediated inflammatory cell recruitment that requires Mac-1-integrin. The EMBO journal 26, 1129–1139 (2007).

40. K. Podar et al., Vascular endothelial growth factor-induced migration of multiple myeloma cells is associated with β1 integrin-and phosphatidylinositol 3-kinase-dependent PKCα activation. Journal of biological chemistry 277, 7875–7881 (2002).

41. R. Bueno et al., Comprehensive genomic analysis of malignant pleural mesothelioma identifies recurrent mutations, gene fusions and splicing alterations. Nature genetics 47, 407 (2015).

42. B. C. Christensen et al., Differentiation of lung adenocarcinoma, pleural mesothelioma, and nonmalignant pulmonary tissues using DNA methylation profiles. Cancer research 69, 6315–6321 (2009).

43. S. Wang et al., Urokinase-type plasminogen activator receptor promotes proliferation and invasion with reduced cisplatin sensitivity in malignant mesothelioma. Oncotarget 7, 69565 (2016).

44. L. A. Marek et al., Nonamplified FGFR1 is a growth driver in malignant pleural mesothelioma. Molecular cancer research 12, 1460–1469 (2014).

45. T. R. Wilson, D. Y. Lee, L. Berry, D. S. Shames, J. Settleman, Neuregulin-1-mediated autocrine signaling underlies sensitivity to HER2 kinase inhibitors in a subset of human cancers. Cancer cell 20, 158–172 (2011).

46. R. Kaarteenaho-Wiik, Y. Soini, R. Pöllänen, P. Pääkkö, V. Kinnula, Over-expression of tenascin-C in malignant pleural mesothelioma. Histopathology 42, 280–291 (2003).

47. C.-C. Lin et al., Malignant pleural effusion cells show aberrant glucose metabolism gene expression. European Respiratory Journal 37, 1453–1465 (2011).

48. S. Crispi et al., Global gene expression profiling of human pleural mesotheliomas: identification of matrix metalloproteinase 14 (MMP-14) as potential tumour target. PLoS One 4, e7016 (2009).

49. G. Consortium, The Genotype-Tissue Expression (GTEx) pilot analysis: Multitissue gene regulation in humans. Science 348, 648–660 (2015).

50. M. Uhlen et al., A pathology atlas of the human cancer transcriptome. Science 357, eaan2507 (2017).

51. M. Horie, A. Saito, M. Ohshima, H. I. Suzuki, T. Nagase, YAP and TAZ modulate cell phenotype in a subset of small cell lung cancer. Cancer science 107, 1755–1766 (2016).

52. L. Strizzi et al., Vascular endothelial growth factor is an autocrine growth factor in human malignant mesothelioma. The Journal of pathology 193, 468–475 (2001).

53. T. Seto et al., Prognostic value of expression of vascular endothelial growth factor and its flt-1 and KDR receptors in stage I non-small-cell lung cancer. Lung cancer 53, 91–96 (2006).

54. N. Yanamala et al., Characterization of pulmonary responses in mice to asbestos/asbestiform fibers using gene expression profiles. Journal of Toxicology and Environmental Health, Part A 81, 60–79 (2018).

55. A. Kramer, J. Green, J. Pollard, Jr., S. Tugendreich, Causal analysis approaches in Ingenuity Pathway Analysis. Bioinformatics 30, 523–530 (2014).

56. J. LoPiccolo, C. A. Granville, J. J. Gills, P. A. Dennis, Targeting Akt in cancer therapy. Anti-cancer drugs 18, 861–874 (2007).

57. K. B. Karunakaran, S. Chaparala, M. K. Ganapathiraju, Potentially repurposable drugs for schizophrenia identified from its interactome. bioRxiv, (2018).

58. A. Chattopadhyay, M. K. Ganapathiraju, Demonstration Study: A Protocol to Combine Online Tools and Databases for Identifying Potentially Repurposable Drugs. Data 2, 15 (2017).

59. I. Kupershmidt et al., Ontology-based meta-analysis of global collections of high-throughput public data. PLoS One 5, (2010).

60. M. Suraokar et al., Expression profiling stratifies mesothelioma tumors and signifies deregulation of spindle checkpoint pathway and microtubule network with therapeutic implications. Annals of Oncology 25, 1184–1192 (2014).

61. H. L. Kindler, J. P. van Meerbeeck, in Seminars in oncology. (Elsevier, 2002), vol. 29, pp. 70–76.

62. N. J. Vogelzang et al., Phase III study of pemetrexed in combination with cisplatin versus cisplatin alone in patients with malignant pleural mesothelioma. Journal of clinical oncology 21, 2636–2644 (2003).

63. A. Scherpereel et al. (American Society of Clinical Oncology, 2017).

64. S. Puhalla, A. Brufsky, Ixabepilone: a new chemotherapeutic option for refractory metastatic breast cancer. Biologics: targets & therapy 2, 505 (2008).

65. M. O. Jamil, M. S. Jerome, D. Miley, K. S. Selander, F. Robert, A pilot study of zoledronic acid in the treatment of patients with advanced malignant pleural mesothelioma. Lung Cancer: Targets and Therapy 8, 39 (2017).

66. S. Okamoto et al., Zoledronic acid produces antitumor effects on mesothelioma through apoptosis and S-phase arrest in p53-independent and Ras prenylation-independent manners. Journal of Thoracic Oncology 7, 873–882 (2012).

67. M.-L. Hartman, J. M. Esposito, B. Y. Yeap, D. J. Sugarbaker, Combined treatment with cisplatin and sirolimus to enhance cell death in human mesothelioma. The Journal of thoracic and cardiovascular surgery 139, 1233–1240 (2010).

68. M. Tomasetti, N. Gellert, A. Procopio, J. Neuzil, A vitamin E analogue suppresses malignant mesothelioma in a preclinical model: a future drug against a fatal neoplastic disease? International journal of cancer 109, 641–642 (2004).

69. J. Kovarova et al., Mitochondrial targeting of α-tocopheryl succinate enhances its anti-mesothelioma efficacy. Redox Report 19, 16–25 (2014).

70. A. Sato, N. Virgona, Y. Sekine, T. Yano, The evidence to date: A redox-inactive analogue of tocotrienol as a new anti-mesothelioma agent. (2016).

71. A. Kotsakis et al., A multicentre phase II trial of cabazitaxel in patients with advanced non-small-cell lung cancer progressing after docetaxel-based chemotherapy. British journal of cancer 115, 784 (2016).

72. L. M. Krug, R. T. Heelan, M. G. Kris, E. Venkatraman, F. Sirotnak, Phase II trial of pralatrexate (10-propargyl-10-deazaaminopterin, PDX) in patients with unresectable malignant pleural mesothelioma. Journal of Thoracic Oncology 2, 317–320 (2007).

73. V. Karantza, Keratins in health and cancer: more than mere epithelial cell markers. Oncogene 30, 127 (2011).

74. O. D. Røe et al., Molecular resistance fingerprint of pemetrexed and platinum in a long-term survivor of mesothelioma. PloS one 7, e40521 (2012).

75. M. Miettinen, J. Limon, A. Niezabitowski, J. Lasota, Calretinin and other mesothelioma markers in synovial sarcoma: analysis of antigenic similarities and differences with malignant mesothelioma. The American journal of surgical pathology 25, 610–617 (2001).

76. A.-R. Choi, J.-H. Kim, Y. H. Woo, H. S. Kim, S. Yoon, Anti-malarial drugs primaquine and chloroquine have different sensitization effects with anti-mitotic drugs in resistant cancer cells. Anticancer research 36, 1641–1648 (2016).

77. T. Kimura et al., Comparisons between the beneficial effects of different sulphonylurea treatments on ischemia-induced retinal neovascularization. Free Radical Biology and Medicine 43, 454–461 (2007).

78. A. Yasumitsu et al., Clinical significance of serum vascular endothelial growth factor in malignant pleural mesothelioma. Journal of Thoracic Oncology 5, 479–483 (2010).

79. N. Hirayama et al., Pleural effusion VEGF levels as a prognostic factor of malignant pleural mesothelioma. Respiratory medicine 105, 137–142 (2011).

80. G. Pasello, L. Urso, P. Conte, A. Favaretto, Effects of sulfonylureas on tumor growth: a review of the literature. The oncologist 18, 1118–1125 (2013).

81. Y. Zhou, R.-A. F. Gunput, R. J. Pasterkamp, Semaphorin signaling: progress made and promises ahead. Trends in biochemical sciences 33, 161–170 (2008).

82. T. Ito et al., Repulsive axon guidance molecule Sema3A inhibits branching morphogenesis of fetal mouse lung. Mechanisms of development 97, 35–45 (2000).

83. T. Kawakami et al., Neuropilin 1 and neuropilin 2 co-expression is significantly correlated with increased vascularity and poor prognosis in nonsmall cell lung carcinoma. Cancer 95, 2196–2201 (2002).

84. G. Neufeld, O. Kessler, The semaphorins: versatile regulators of tumour progression and tumour angiogenesis. Nature reviews. Cancer 8, 632 (2008).

85. K. J. Beauchemin et al., Temporal dynamics of the developing lung transcriptome in three common inbred strains of laboratory mice reveals multiple stages of postnatal alveolar development. PeerJ 4, e2318 (2016).

86. Y. Okada et al., Genetics of rheumatoid arthritis contributes to biology and drug discovery. Nature 506, 376–381 (2014).

87. E. Oliemuller et al., Phosphorylated tubulin adaptor protein CRMP-2 as prognostic marker and candidate therapeutic target for NSCLC. International journal of cancer 132, 1986–1995 (2013).

88. J. Guo, The roles of β tubulin isotypes in neurons. (The University of Texas Health Science Center at San Antonio, 2011).

89. K.-H. Braunewell, A. J. K. Szanto, Visinin-like proteins (VSNLs): interaction partners and emerging functions in signal transduction of a subfamily of neuronal Ca2+-sensor proteins. Cell and tissue research 335, 301–316 (2009).

90. P. Nymark et al., Integrative analysis of microRNA, mRNA and aCGH data reveals asbestos-and histology-related changes in lung cancer. Genes, Chromosomes and Cancer 50, 585–597 (2011).

91. Y. Sasaki et al., Fyn and Cdk5 mediate semaphorin-3A signaling, which is involved in regulation of dendrite orientation in cerebral cortex. Neuron 35, 907–920 (2002).

92. P. Poulikakos et al., Re-expression of the tumor suppressor NF2/merlin inhibits invasiveness in mesothelioma cells and negatively regulates FAK. Oncogene 25, 5960–5968 (2006).

93. U. Pley, P. Parham, F. M. Brodsky, Clathrin: its role in receptor-mediated vesicular transport and specialized functions in neurons. Critical reviews in biochemistry and molecular biology 28, 431–464 (1993).

94. G. Gallo, P. C. Letourneau, Regulation of growth cone actin filaments by guidance cues. Developmental Neurobiology 58, 92–102 (2004).

95. C.-H. Tsai et al., Over-expression of cofilin-1 suppressed growth and invasion of cancer cells is associated with up-regulation of let-7 microRNA. Biochimica et Biophysica Acta (BBA)-Molecular Basis of Disease 1852, 851–861 (2015).

96. D. Carbonari et al., Angiogenic effect induced by mineral fibres. Toxicology 288, 34–42 (2011).

97. N. Kumagai-Takei et al., Asbestos induces reduction of tumor immunity. Clinical and Developmental Immunology 2011, (2011).

98. S. Sneddon et al., Whole exome sequencing of an asbestos-induced wild-type murine model of malignant mesothelioma. BMC cancer 17, 396 (2017).

99. L. Fang et al., MiR-93 enhances angiogenesis and metastasis by targeting LATS2. Cell cycle 11, 4352-4365 (2012).

100. W. Zhu et al., Expression of miR-29c, miR-93, and miR-429 as potential biomarkers for detection of early stage non-small lung cancer. PloS one 9, e87780 (2014).

101. S. Busacca et al., MicroRNA signature of malignant mesothelioma with potential diagnostic and prognostic implications. American journal of respiratory cell and molecular biology 42, 312–319 (2010).

102. H. W. Hopf et al., Hyperoxia and angiogenesis. Wound repair and regeneration 13, 558–564 (2005).

103. R. M. Ryan et al., in D108. LESSONS IN ANGIOGENESIS ACROSS DEVELOPMENT AND DISEASE. (Am Thoracic Soc, 2013), pp. A5908–A5908.

104. T. Fukamachi, S. Ikeda, H. Saito, M. Tagawa, H. Kobayashi, Expression of acidosis-dependent genes in human cancer nests. Molecular and clinical oncology 2, 1160–1166 (2014).

105. L. R. Pearce et al., Identification of Protor as a novel Rictor-binding component of mTOR complex-2. Biochemical Journal 405, 513–522 (2007).

106. R. Zoncu, D. M. Sabatini, A. Efeyan, mTOR: from growth signal integration to cancer, diabetes and ageing. Nature reviews. Molecular cell biology 12, 21 (2011).

107. S.-Y. Woo et al., PRR5, a novel component of mTOR complex 2, regulates platelet-derived growth factor receptor β expression and signaling. Journal of Biological Chemistry 282, 25604–25612 (2007).

108. R. Baron, M. Kneissel, WNT signaling in bone homeostasis and disease: from human mutations to treatments. Nature medicine 19, 179–192 (2013).

109. J. Chen et al., WNT7B promotes bone formation in part through mTORC1. PLoSgenetics 10, e1004145 (2014).

110. X. Wang et al., Expression of histone deacetylase 3 instructs alveolar type I cell differentiation by regulating a Wnt signaling niche in the lung. Developmental biology 414, 161–169 (2016).

111. H. Liu et al., CUB-domain–containing protein 1 (CDCP1) activates Src to promote melanoma metastasis. Proceedings of the National Academy of Sciences 108, 1379–1384 (2011).

112. R.-B. Yang et al., Identification of a novel family of cell-surface proteins expressed in human vascular endothelium. Journal of Biological Chemistry 277, 46364–46373 (2002).

113. Y.-C. Lin et al., Endothelial SCUBE2 interacts with VEGFR2 and regulates VEGF-induced angiogenesis. Arteriosclerosis, thrombosis, and vascular biology, ATVBAHA. 116.308546 (2016).

114. A. K. Das, P. T. Cohen, D. Barford, The structure of the tetratricopeptide repeats of protein phosphatase 5: implications for TPR-mediated protein–lprotein interactions. The EMBOjournal 17, 1192–1199 (1998).

115. H. Kim, Y. Phung, M. Ho, Changes in global gene expression associated with 3D structure of tumors: an ex vivo matrix-free mesothelioma spheroid model. PLoS One 7, e39556 (2012).

116. M. Zanoni et al., 3D tumor spheroid models for in vitro therapeutic screening: a systematic approach to enhance the biological relevance of data obtained. Scientific reports 6, (2016).

117. A. Pellagatti et al., Deregulated gene expression pathways in myelodysplastic syndrome hematopoietic stem cells. Leukemia 24, 756 (2010).

118. M. G. Vander Heiden, L. C. Cantley, C. B. Thompson, Understanding the Warburg effect: the metabolic requirements of cell proliferation. science 324, 1029–1033 (2009).

119. M. Tarrado-Castellarnau, P. de Atauri, M. Cascante, Oncogenic regulation of tumor metabolic reprogramming. Oncotarget 7, 62726 (2016).

120. K. G. Bulock, G. P. Beardsley, K. S. Anderson, The kinetic mechanism of the human bifunctional enzyme ATIC (5-amino-4-imidazolecarboxamide ribonucleotide transformylase/inosine 5’-monophosphate cyclohydrolase) a surprising lack of substrate channeling. Journal of Biological Chemistry 277, 22168-22174 (2002).

121. M. Çeliktaş et al., Role of CPS1 in cell growth, metabolism, and prognosis in LKB1-inactivated lung adenocarcinoma. Journal of the National Cancer Institute 109, djw231 (2016).

122. A. Nakano, S. Takashima, LKB1 and AMP-activated protein kinase: regulators of cell polarity. Genes to Cells 17, 737–747 (2012).

123. D. C. Lam et al., Oncogenic mutation profiling in new lung cancer and mesothelioma cell lines. OncoTargets and therapy 8, 195 (2015).

124. D. B. Shackelford et al., LKB1 inactivation dictates therapeutic response of non-small cell lung cancer to the metabolism drug phenformin. Cancer cell 23, 143–158 (2013).

125. D. J. Asby et al., AMPK activation via modulation of de novo purine biosynthesis with an inhibitor of ATIC homodimerization. Chemistry & biology 22, 838–848 (2015).

126. J. Kim, G. Yang, Y. Kim, J. Kim, J. Ha, AMPK activators: mechanisms of action and physiological activities. Experimental & molecular medicine 48, e224 (2016).

127. X. Liu et al., The complex genetics of hypoplastic left heart syndrome. Nat Genet 49, 1152–1159 (2017).

128. M. R. Junttila et al., CIP2A inhibits PP2A in human malignancies. Cell 130, 51–62 (2007).

129. W. Li et al., CIP2A is overexpressed in gastric cancer and its depletion leads to impaired clonogenicity, senescence, or differentiation of tumor cells. Clinical cancer research 14, 3722–3728 (2008).

130. M. Toyoshima et al., Functional genomics identifies therapeutic targets for MYC-driven cancer. Proceedings of the National Academy of Sciences 109, 9545–9550 (2012).

131. X. Zhang, D. Yee, Tyrosine kinase signalling in breast cancer: insulin-like growth factors and their receptors in breast cancer. Breast Cancer Research 2, 170 (2000).

132. C. D. Hoang et al., Selective activation of insulin receptor substrate-1 and-2 in pleural mesothelioma cells. Cancer research 64, 7479–7485 (2004).

133. F. Hasteh, G. Y. Lin, N. Weidner, C. W. Michael, The use of immunohistochemistry to distinguish reactive mesothelial cells from malignant mesothelioma in cytologic effusions. Cancer cytopathology 118, 90–96 (2010).

134. Y. Ohyama et al., Modulation of matrix mineralization by Vwc2-like protein and its novel splicing isoforms. Biochemical and biophysical research communications 418, 12–16 (2012).

135. A. Miyake et al., Brorin is required for neurogenesis, gliogenesis, and commissural axon guidance in the zebrafish forebrain. PloS one 12, e0176036 (2017).

136. E. L. Huttlin et al., The BioPlex network: a systematic exploration of the human interactome. Cell 162, 425–440 (2015).

137. D. A. Fennell, G. Gaudino, K. J. O’byrne, L. Mutti, J. Van Meerbeeck, Advances in the systemic therapy of malignant pleural mesothelioma. Nature Reviews. Clinical Oncology 5, 136 (2008).

138. K. Donaldson, F. A. Murphy, R. Duffin, C. A. Poland, Asbestos, carbon nanotubes and the pleural mesothelium: a review of the hypothesis regarding the role of long fibre retention in the parietal pleura, inflammation and mesothelioma. Particle and fibre toxicology 7, 5 (2010).

139. B. Lu et al., Novel role of PKR in inflammasome activation and HMGB1 release. Nature 488, 670 (2012).

140. D. Musumeci, G. N. Roviello, D. Montesarchio, An overview on HMGB1 inhibitors as potential therapeutic agents in HMGB1-related pathologies. Pharmacology & therapeutics 141, 347–357 (2014).

141. F. Biscetti et al., High-mobility group box-1 protein promotes angiogenesis after peripheral ischemia in diabetic mice through a VEGF-dependent mechanism. Diabetes 59, 1496–1505 (2010).

142. R. Kang, Q. Zhang, H. J. Zeh, M. T. Lotze, D. Tang, HMGB1 in cancer: good, bad, or both? Clinical cancer research 19, 4046–4057 (2013).

143. J. Van Beijnum et al., Tumor angiogenesis is enforced by autocrine regulation of high-mobility group box 1. Oncogene 32, 363 (2013).

144. D. Tang, R. Kang, H. J. Zeh, M. T. Lotze, High-mobility group box 1 and cancer. Biochimica et Biophysica Acta (BBA)-Gene Regulatory Mechanisms 1799, 131–140 (2010).

145. R. Grose, C. Dickson, Fibroblast growth factor signaling in tumorigenesis. Cytokine & growth factor reviews 16, 179–186 (2005).

146. M. L. Galisteo, J. Chernoff, Y.-C. Su, E. Y. Skolnik, J. Schlessinger, The adaptor protein Nck links receptor tyrosine kinases with the serine-threonine kinase Pak1. Journal of Biological Chemistry 271, 20997–21000 (1996).

147. J.-i. Nitadori et al., Immunohistochemical differential diagnosis between large cell neuroendocrine carcinoma and small cell carcinoma by tissue microarray analysis with a large antibody panel. American journal of clinical pathology 125, 682–692 (2006).

148. M. Taipale, D. F. Jarosz, S. Lindquist, HSP90 at the hub of protein homeostasis: emerging mechanistic insights. Nature reviews. Molecular cell biology 11, 515 (2010).

149. W. Huang et al., HMGB1 increases permeability of the endothelial cell monolayer via RAGE and Src family tyrosine kinase pathways. Inflammation 35, 350–362 (2012).

150. C. Chen, T. Lou, Hypoxia inducible factors in hepatocellular carcinoma. Oncotarget 8, 46691 (2017).

151. M. B. Marron, D. P. Hughes, M. D. Edge, C. L. Forder, N. P. Brindle, Evidence for heterotypic interaction between the receptor tyrosine kinases TIE-1 and TIE-2. Journal of Biological Chemistry 275, 39741–39746 (2000).

152. P. Sawma et al., Evidence for new homotypic and heterotypic interactions between transmembrane helices of proteins involved in receptor tyrosine kinase and neuropilin signaling. Journal of molecular biology 426, 4099–4111 (2014).

153. A. A. Lanahan et al., VEGF receptor 2 endocytic trafficking regulates arterial morphogenesis. Developmental cell 18, 713–724 (2010).

154. N. Volodko, M. Gordon, M. Salla, H. A. Ghazaleh, S. Baksh, RASSF tumor suppressor gene family: biological functions and regulation. FEBS letters 588, 2671–2684 (2014).

155. R. A. Kahn, M. P. East, J. W. Francis, in Ras Superfamily Small G Proteins: Biology and Mechanisms 2. (Springer, 2014), pp. 215–251.

156. R. W. Cole, J. G. Ault, J. H. Hayden, C. L. Rieder, Crocidolite asbestos fibers undergo size-dependent microtubule-mediated transport after endocytosis in vertebrate lung epithelial cells. Cancer research 51, 4942–4947 (1991).

157. T. L. Whiteside, in Seminars in cancer biology. (Elsevier, 2006), vol. 16, pp. 3–15.

158. H. Batra, V. B. Antony, Pleural mesothelial cells in pleural and lung diseases. Journal of thoracic disease 7, 964 (2015).

159. J. Rautela et al., Loss of host type-I IFN signaling accelerates metastasis and impairs NK-cell antitumor function in multiple models of breast cancer. Cancer immunology research 3, 1207–1217 (2015).

160. J. B. Swann et al., Type I IFN contributes to NK cell homeostasis, activation, and antitumor function. The Journal of Immunology 178, 7540–7549 (2007).

161. S. Y. Al Omar, E. Marshall, D. Middleton, S. E. Christmas, Increased killer immunoglobulin-like receptor expression and functional defects in natural killer cells in lung cancer. Immunology 133, 94–104 (2011).

162. E.-C. Shin, K. S. Choi, S. J. Kim, J.-S. Shin, Modulation of the Surface Expression of CD158 Killer Cell Ig-like Receptor by Interleukin-2 and Transforming Growth. Yonsei medical journal 45, 510–514 (2004).

163. S. J. Parsons, J. T. Parsons, Src family kinases, key regulators of signal transduction. Oncogene 23, 7906 (2004).

164. M. T. Goswami et al., Role and regulation of coordinately expressed de novo purine biosynthetic enzymes PPAT and PAICS in lung cancer. Oncotarget 6, 23445 (2015).

165. G. Luengo-Gil et al., Antithrombin controls tumor migration, invasion and angiogenesis by inhibition of enteropeptidase. Scientific reports 6, (2016).

166. J. Hou et al., N-Myc-interacting protein (NMI) negatively regulates epithelial-mesenchymal transition by inhibiting the acetylation of NF-_K_B/p65. Cancer letters 376, 22–33 (2016).

167. M.-M. Tsai et al., MicroRNA-26b inhibits tumor metastasis by targeting the KPNA2/c-jun pathway in human gastric cancer. Oncotarget 7, 39511 (2016).

168. T. S. Khatlani et al., c-Jun N-terminal kinase is activated in non-small-cell lung cancer and promotes neoplastic transformation in human bronchial epithelial cells. Oncogene 26, 2658 (2007).

169. R. J. Buchsbaum, B. A. Connolly, L. A. Feig, Interaction of Rac exchange factors Tiam1 and Ras-GRF1 with a scaffold for the p38 mitogen-activated protein kinase cascade. Molecular and cellular biology 22, 4073–4085 (2002).

170. N. Welchering et al., Dexmedetomidine and fentanyl exhibit temperature dependent effects on human respiratory cilia. Frontiers in pediatrics 3, 7 (2015).

171. P. Sathe et al., Innate immunodeficiency following genetic ablation of Mcl1 in natural killer cells. Nature communications 5, 4539 (2014).

172. S. G. Crone et al., microRNA-146a inhibits G protein-coupled receptor-mediated activation of NF-κB by targeting CARD10 and COPS8 in gastric cancer. Molecular cancer 11, 71 (2012).

173. S. W. Bailey, J. E. Ayling, The extremely slow and variable activity of dihydrofolate reductase in human liver and its implications for high folic acid intake. Proceedings of the National Academy of Sciences 106, 15424–15429 (2009).

174. N. Hagner, M. Joerger, Cancer chemotherapy: targeting folic acid synthesis. Cancer management and research 2, 293 (2010).

175. N. H. Heintz, Y. M. Janssen-Heininger, B. T. Mossman, Asbestos, lung cancers, and mesotheliomas: from molecular approaches to targeting tumor survival pathways. American journal of respiratory cell and molecular biology 42, 133–139 (2010).

176. S. F. Tavazoie et al., Endogenous human microRNAs that suppress breast cancer metastasis. nature 451, 147 (2008).

177. M. Tagawa, Y. Tada, H. Shimada, K. Hiroshima, Gene therapy for malignant mesothelioma: current prospects and challenges. Cancer gene therapy 20, 150 (2013).

178. A. Krämer, J. Green, J. Pollard Jr, S. Tugendreich, Causal analysis approaches in ingenuity pathway analysis. Bioinformatics 30, 523–530 (2013).

179. P. Shannon et al., Cytoscape: a software environment for integrated models of biomolecular interaction networks. Genome research 13, 2498–2504 (2003).

180. Y. Tanaka et al., Structural implications for heavy metal-induced reversible assembly and aggregation of a protein: the case of Pyrococcus horikoshii CutA 1. FEBS letters 556, 167–174 (2004).

181. P. J. Thul et al., A subcellular map of the human proteome. Science 356, eaal3321 (2017).

182. M. Uhlén et al., Tissue-based map of the human proteome. Science 347, 1260419 (2015).

183. F. M. Johnson et al., Phase II study of dasatinib in patients with advanced non–small-cell lung cancer. Journal of clinical oncology 28, 4609–4615 (2010).

184. A. S. Tsao et al., Biomarker-Integrated Neoadjuvant Dasatinib Trial in Resectable Malignant Pleural Mesothelioma. Journal of Thoracic Oncology, (2017).

185. A. M Comer, K. L Goa, Docetaxel. A review of its use in non-small cell lung cancer. (2000), vol. 17, pp. 53–80.

186. C. P. Belani et al., Docetaxel for malignant mesothelioma: phase II study of the Eastern Cooperative Oncology Group. Clinical lung cancer 6, 43–47 (2004).

187. M. Ralli et al., Docetaxel plus gemcitabine as first-line treatment in malignant pleural mesothelioma: a single institution phase II study. Anticancer research 29, 3441–3444 (2009).

188. I. Tourkantonis et al., Phase II study of gemcitabine plus docetaxel as second-line treatment in malignant pleural mesothelioma: a single institution study. American journal of clinical oncology 34, 38–42 (2011).

189. C. Manegold, Gemcitabine (Gemzar®) in non-small cell lung cancer. Expert review of anticancer therapy 4, 345–360 (2004).

190. J. Malhotra, S. K. Jabbour, J. Aisner, Current state of immunotherapy for non-small cell lung cancer. Translational lung cancer research 6, 196 (2017).

191. D. R. Spigel et al., Phase II trial of ixabepilone and carboplatin with or without bevacizumab in patients with previously untreated advanced non–small-cell lung cancer. Lung Cancer 78, 70–75 (2012).

192. N. Altorki et al., Phase II proof-of-concept study of pazopanib monotherapy in treatment-naive patients with stage I/II resectable non–small-cell lung cancer. Journal of Clinical Oncology 28, 3131–3137 (2010).

193. B. I. Hiddinga, C. Rolfo, J. P. van Meerbeeck, Mesothelioma treatment: Are we on target? A review. Journal of advanced research 6, 319–330 (2015).

194. G. V. Scagliotti et al., Phase III study comparing cisplatin plus gemcitabine with cisplatin plus pemetrexed in chemotherapy-naive patients with advanced-stage non–small-cell lung cancer. Journal of clinical oncology 26, 3543–3551 (2008).

195. T. Shimizu et al., Association between expression of thymidylate synthase, dihydrofolate reductase, and glycinamide ribonucleotide formyltransferase and efficacy of pemetrexed in advanced non-small cell lung cancer. Anticancer research 32, 4589–4596 (2012).

196. A. M. Giammarioli et al., Pyrimethamine induces apoptosis of melanoma cells via a caspase and cathepsin double-edged mechanism. Cancer research 68, 5291–5300 (2008).

197. M. W. Khan et al., The STAT3 inhibitor pyrimethamine displays anti-cancer and immune stimulatory effects in murine models of breast cancer. Cancer Immunology, Immunotherapy 67, 13–23 (2018).

198. W. E. Grose, G. P. Bodey, V. Rodriguez, Sulfamethoxazole-trimethoprim for infections in cancer patients. JAMA 237, 352–354 (1977).

199. G. P. Bodey, W. E. Grose, M. J. Keating, Use of trimethoprim-sulfamethoxazole for treatment of infections in patients with cancer. Reviews of infectious diseases 4, 579–585 (1982).

200. S.-H. Lu et al., Identifying cancer origin using circulating tumor cells. Cancer biology & therapy 17, 430-438 (2016).

201. R. Camilo, V. L. Capelozzi, S. A. C. Siqueira, F. D. C. Bernardi, Expression of p63, keratin 5/6, keratin 7, and surfactant-A in non–small cell lung carcinomas. Human pathology 37, 542–546 (2006).

202. N. G. Ordóñez, Value of cytokeratin 5/6 immunostaining in distinguishing epithelial mesothelioma of the pleura from lung adenocarcinoma. The American journal of surgical pathology 22, 1215–1221 (1998).

203. C. Heard, B. Monk, A. Modley, Binding of primaquine to epidermal membranes and keratin. International journal of pharmaceutics 257, 237–244 (2003).

204. T. Kimura et al., The antimalarial drugs chloroquine and primaquine inhibit pyridoxal kinase, an essential enzyme for vitamin B6 production. FEBS letters 588, 3673–3676 (2014).

205. L. G. Basso, R. Z. Rodrigues, R. M. Naal, A. J. Costa-Filho, Effects of the antimalarial drug primaquine on the dynamic structure of lipid model membranes. Biochimica et Biophysica Acta (BBA)-Biomembranes 1808, 55–64 (2011).

206. G. Gakhar, T. Ohira, S. Battina, D. H. Hua, T. A. Nguyen. (AACR, 2007).

207. S.-H. I. Ou et al., SWOG S0722: phase II study of mTOR inhibitor everolimus (RAD001) in advanced malignant pleural mesothelioma (MPM). Journal of Thoracic Oncology 10, 387–391 (2015).

208. J.-C. Mamputu, G. Renier, Advanced glycation end products increase, through a protein kinase C-dependent pathway, vascular endothelial growth factor expression in retinal endothelial cells: inhibitory effect of gliclazide. Journal of Diabetes and its Complications 16, 284–293 (2002).

